# Mathematical modelling of macrophage and natural killer cell immune response during early stages of peritoneal endometriosis lesion onset

**DOI:** 10.1101/2025.01.20.633967

**Authors:** Claire M. Miller, Domenic P.J. Germano, Alicia M. Chenoweth, Sarah Holdsworth-Carson

## Abstract

The immune system is hypothesised to contribute to the onset of endometriosis lesions. However, the precise mechanisms underlying its role are not yet known. We introduce a novel compartmental model that describes the interactions between innate immune cells, specifically macrophages and natural killer cells, and endometrial cells, occurring within the peritoneal fluid during the early stages of (superficial peritoneal) endometriosis lesion onset. Our study focuses on retrograde influx, immune detection, and immune clearance. Results show an increased influx of endometrial cells into peritoneal fluid correlates with heightened pro-inflammatory macrophage activation, but does not lead to an increase in disease. We compare the system’s response to changes in immune cytotoxicity and ability to detect ectopic endometrial cells. We predict that reduced cytotoxicity is a key driver of disease. These findings align with the increased immune activation observed clinically. Lastly, we predict that an individual can transition to a diseased state following a reduction in immune system cytotoxicity and/or reduced ability to detect ectopic cells. Due to hysteresis, a significant improvement is then required to restore an individual to the disease-free state. This work provides a valuable framework to explore hypotheses of endometriosis lesion onset and assist in understanding of the disease.

## 1 Background

Endometriosis is a chronic gynaecological condition that affects approximately one in nine women [1]. The disease is characterised by the presence of endometrial-like tissue (tissue similar to the lining of the uterus) growing in lesions outside of the uterus. Endometriosis results in a variety of symptoms, including chronic pain and fertility issues [2, 3].

Despite its high prevalence, the mechanisms behind the onset of endometriosis remain an open question [1, 2]. One hypothesis is Sampson’s theory of retrograde menstruation [2–4]. Under this hypothesis, some menstrual debris, which includes endometrial stromal and epithelial cells as well as immune cells, is cleared from the uterus via the fallopian tubes, rather than through the cervix, during menstruation. After exiting the fallopian tubes, endometrial cells and other menstrual debris collect in the peritoneal fluid (4–16mL [5, 6]) within the peritoneal cavity. Endometrial cells are carried by this fluid and some of these cells may attach to internal tissues, invade the tissue, and develop into lesions.

However, studies have found the prevalence of people with retrograde menstruation is greater than the prevalence of endometriosis [7], indicating that the retrograde menstruation theory alone is insufficient to explain disease onset. Supporting this, a recent review has questioned whether previous evidence is sufficient to confirm differences in frequency and volume of retrograde menstruation in endometriosis patients [8]. This leads to two key hypotheses: in endometriosis patients, a normally functioning immune system is overcome by an increased concentration of endometrial cells from menstrual debris, or (B) there exists an additional dysfunction of the immune system in those that experience disease. These hypotheses are still open questions for endometriosis pathophysiology [2].

In support of Hypothesis A, studies have shown a decreased proportion of cells undergoing apoptosis within eutopic endometrium of endometriosis patients [9], indicating an increased number of viable endometrial cells with the potential to form lesions enter peritoneal fluid. Endometriosis has also been associated with shorter menstrual cycle length, increased menstrual flow duration, and heavy menstrual flow volume [10–12], which all indicate increased influx into the peritoneal cavity. However, other studies have shown no significant difference in clonogenicity or concentrations of endometrial mesenchymal stromal and epithelial stem cells in both peritoneal fluid and menstrual blood between endometriosis patients and controls, although much higher variation was observed in the endometriosis patients [13].

Ectopic endometrial cells in the peritoneal cavity should be cleared by the immune system, preventing lesion formation. Consequently, a dysfunction in the ability of the immune system to detect or clear the cells could be involved in disease onset (Hypothesis B) [14]. Several studies have shown differences in immune cell profiles in the peritoneal fluid of endometriosis patients compared to controls [3, 15–18]. Due to the complexity of the immune system, it is difficult from observation alone to identify which differences are causes of immune dysfunction, a consequence of immune dysfunction, or simply a consequence of the presence of the endometrial cells. Mathematical modelling enables us to understand the relationship between hypothesised immune dysfunctions and emergent immune system profiles, for example viral infections [19] and cancer growth [20].

The early immune response is dominated by the innate immune system. Macrophages and natural killer (NK) cells are two innate immune cell types commonly implicated in the endometriosis literature. Macrophages contribute to the inflammatory environment of the peritoneal fluid [21, 22], and are involved in the clearance of endometrial cells through phagocytosis and detection of apoptotic markers [23, 24]. NK cells help clear endometrial cells via their strong cytotoxic activities [2, 3, 25].

Macrophages in the peritoneal cavity originate from endometrium, circulating monocytes, and tissue-resident macrophages (both monocyte-derived and embryonic-derived). Different macrophage phenotypes have been hypothesised to either contribute to lesion growth (endometrial), or protect against lesion establishment (monocyte-derived tissue-resident) [26]. Traditionally, macrophage activation states have been categorised as *pro-inflammatory* (also known as M1-type or classically activated macrophages) or *pro-repair* (also known as M2-type or alternately activated macrophages). This binary classification is, for this study, a useful simplification of a macrophage activation spectrum [27, 28]. We define pro-inflammatory macrophages as macrophages that secrete inflammatory cytokines and likely contribute to maintaining a level of chronic inflammation that is associated with endometriosis [2, 22]. We define pro-repair macrophages as macrophages that are anti-inflammatory and are known to promote lesion growth, through upregulation of angiogenic factors and endometrial cell proliferation [2, 29]. For convenience, we refer to pro-inflammatory macrophages as *M1-type* and pro-repair macrophages as *M2-type* in this paper.

Studies exploring macrophage activation in the peritoneal fluid of endometriosis patients consistently find an increased proportion of macrophages expressing markers of activation [15–18]. These studies have shown a significant increase in the proportion of cells expressing M2-type markers [16–18]. In eutopic endometrium, increased M2-type activation has been observed to be associated with late-stage disease [30]. Ectopic endometrial stromal cells in co-culture have also been shown to induce macrophage secretion of cytokines associated with the promotion of M2-type activation [22, 31, 32]. Increases in macrophages expressing M1-type markers in peritoneal fluid have also been observed, but to a smaller (or non-significant) degree compared to M2-type [16–18]. However, observations of increased pro-inflammatory cytokine expression in peritoneal fluid [3, 18, 21], and increased secretion of IL-1 [3, 21] and cytotoxicity of peritoneal macrophages (from peritoneal fluid) in early-stage disease [33] are all associated with an M1-type activation. Cytotoxicity of peritoneal macrophages (from peritoneal fluid) has also been shown to decrease in late-stage (stage III/IV) disease [33]. Together, this suggests M1-type activation may increase in early but not late stage disease, reflecting an evolving disease environment that could explain the high variability in M1-type activation across patients. Observations of decreased peritoneal macrophage cytotoxicity against both eutopic and ectopic endometrial cells compared to peripheral macrophages in endometriosis patients [34] could also indicate immune function changes as a response to persistent exposure to an altered immune environment, rather than a pre-existing dysfunction. A potential contributor to the inflammatory peritoneal environment of endometriosis patients is the increased M1-type activation in eutopic endometrium which has been observed in endometriosis patients [35]. Endometrial stromal fibroblasts from eutopic endometrium of endometriosis patients also display pro-inflammatory profiles and progesterone resistance [36]. Additionally, a correlation between activated fibroblasts and macrophage populations has been observed in lesion microenvironments [37]. These observations point to a complex relationship between stromal fibroblasts and immune cells in both cycling endometrium and lesion growth.

Macrophages mediate the clearance of cells through the detection of apoptotic markers and phagocytosis [38], which have been shown to be dysregulated in endometriosis lesion cells [39–41]. Several studies have established that ectopic endometrial tissue does not display the same cyclic changes in apoptotic markers as eutopic tissue [42, 43], has a decreased level of apoptosis compared to eutopic endometrium in the same patient [44], and increased anti-apoptotic and decreased pro-apoptotic marker expression compared to eutopic endometrium [42, 43, 45]. Decreased estrogen receptor expression is associated with upregulation of anti-apoptotic markers [43]. In endometriosis patients, decreased [46] and out of phase [47] variation in estrogen receptors on ectopic cells have been observed over the menstrual cycle. In eutopic endometrium, a reduction in spontaneous apoptosis has also been observed [45]. These observations indicate a reduction in expression of apoptotic markers on these cells, resulting in a decrease in detection of these cells by the immune system (Hypothesis B1).

Natural killer (NK) cells are categorised into two types: CD56^bright^ (also known as CD56^bright^CD16^−^) and CD56^dim^ (also known as CD56^dim^CD16^+^). CD56^bright^ cells have high cytokine production while CD56^dim^ cells have higher cytotoxicity [14]. In the endometrium, the majority of NK cells are CD56^bright^ [2]. However, the NK cell profiles in peritoneal fluid during normal function are not fully described. There is mixed evidence on the activation of NK cells in peritoneal fluid of endometriosis patients. After activation by monocytes, NK cells release IFN-γ and TNF-α, the latter of which is consistently raised in endometriosis patients [14]. However, the activity of NK cells has been shown to be decreased in endometriosis patients compared to controls [25]. Peritoneal fluid from endometriosis patients has been shown to reduce cytotoxicity in NK cells *in vitro* [25, 48]. Some studies show IFN-γ induces apoptosis in eutopic but not ectopic endometrial cells [14]. This leads to the hypothesis of reduced clearance of the ectopic endometrial cells by NK cells (Hypothesis B2).

In this paper, we describe a novel compartmental mathematical model of the immune response to endometrial cells in peritoneal fluid. Endometriosis lesions are defined by depth of infiltration (superficial or deep) and location (peritoneal or ovarian) [47]. Our model focuses on the early stages of superficial peritoneal endometriosis lesion onset and immune response, and consequently we only consider innate immune cells. We model the effects of NK cells and macrophages, as these are the cells most commonly hypothesised to be involved in endometriosis onset, as detailed above. We focus on the two different immune cell functions and their potential role in disease, as opposed to the roles of different endometrial cell types. Using this model, we investigate two questions around endometriosis onset:

1. How does the peritoneal fluid immune cell profile change when the amount of retrograde influx is varied, and can increased influx explain observed immune cell profiles in endometriosis patients and disease onset?
2. Which immune system disorder: reduced detection or reduced clearance of endometrial cells by the immune system, is most associated with disease and is consistent with the observed immune cell profiles?

## 2 Methods

In this section we describe the model assumptions governing the interaction between the macrophages, NK cells, and endometrial cells, and define three emergent attachment states which dictate the disease-free and diseased states.

### 2.1 Model

We model the interaction between macrophages *(M*), natural killer cells (*K*), and endometrial (stromal and epithelial) cells (*E*) in peritoneal fluid. Each cell type has the following activation states:

- Macrophages are **resting** (*M*_0_), pro-inflammatory/**M1-type** (*M*_1_) or pro-repair/**M2-type** (*M*_2_).
- Natural killer cells are **resting** (*K*_0_) or **activated** (*K*_*A*_).
- Endometrial cells are **eutopic** (*E*_0_), **in peritoneal fluid** (*E*_*F*_) or **attached** (*E*_*A*_).

We consider attached endometrial cells to be early lesion cells. A diagram of the system model is given in Fig. 1(a) and explanations of each of the cell state transitions and cell interactions is given below.

**Figure 1:**
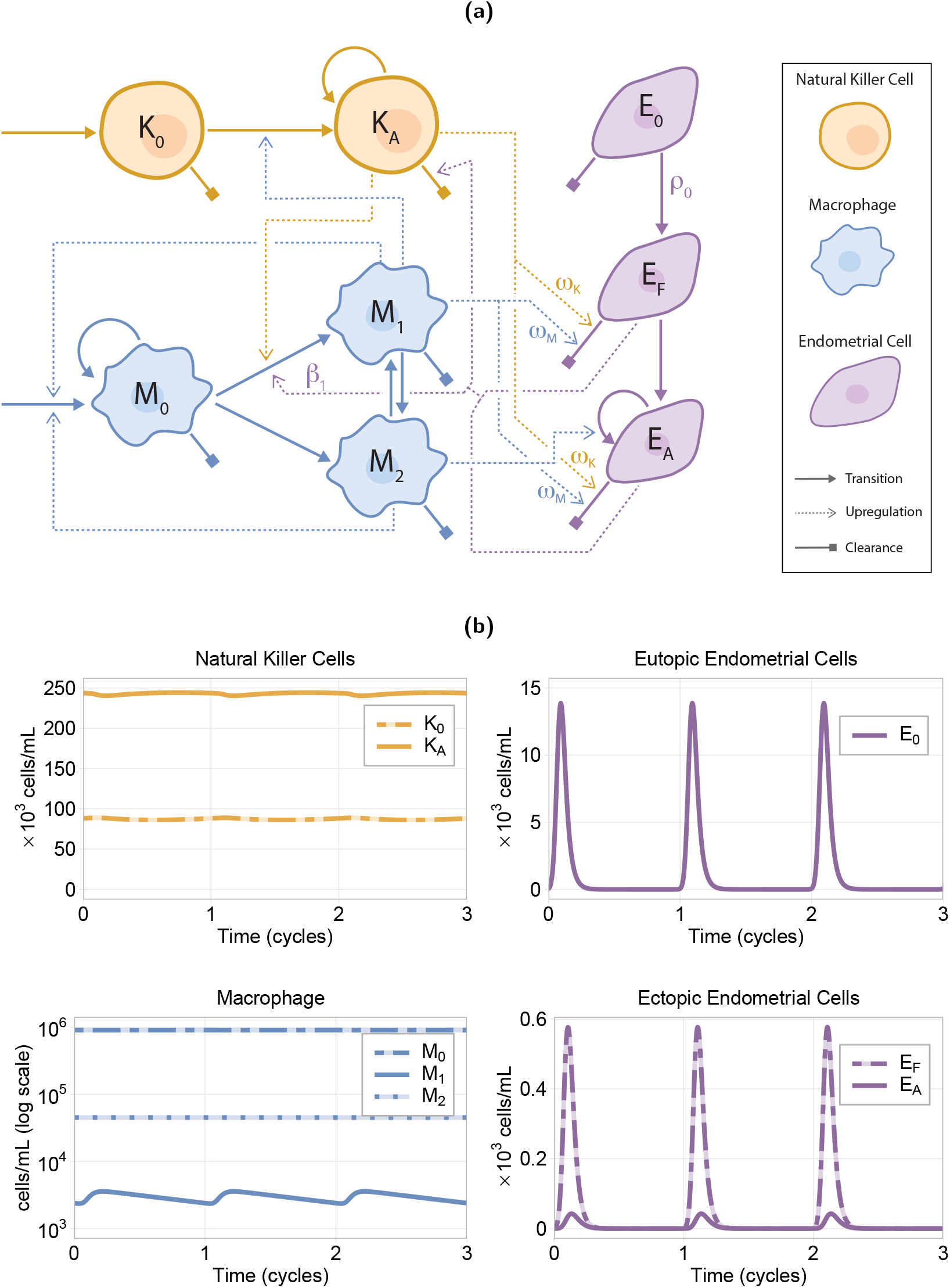
(a) Model diagram with resting (*K*_0_) and activated (*K*_*A*_) natural killer cells (orange); resting (*M*_0_), M1-type (*M*_1_), and M2-type (*M*_2_) macrophages (blue); and eutopic (*E*_0_), in fluid (*E*_*F*_), and attached (*E*_*A*_) endometrial cells (purple). Arrowed solid lines represent cell influx, transition, and proliferation; square arrows represent the cell clearance; and dashed lines represent upregulation of a process. The rate parameters of interest for this study are indicated, where *ω*_*K*_ = *γω* and *ω*_*M*_ = (1 − *γ*)*ω*. (b) Endometrial and immune cell dynamics over three menstrual cycles (on average 28 days) using the default parameter values (see Supplementary S2). The peak in the eutopic endometrial cells, *E*_0_ (top right of (b)), is the menstrual phase of the cycle and these peaks are reflected in the ectopic endometrial cells, *E*_*F*_ and *E*_*A*_ (bottom right of (b)). Similar oscillations can also be seen in the immune system cells (left column of (b)). We see all endometrial cells are cleared before the end of each cycle.

#### 2.1.1 Macrophages

We denote macrophage as resting, *M*_0_, which includes both monocytes from circulation and tissue-resident macrophage. These can be activated to either a pro-inflammatory M1-type, *M*_1_, or pro-repair M2-type, *M*_2_. This binary classification of activated macrophages reflects the average behaviours of a macrophage population activated across a pro-inflammatory to pro-repair spectrum.

##### Macrophage turnover

Macrophage population growth occurs due to: a) recruitment from the blood stream (circulating monocytes); or b) resting macrophage proliferation (tissue-resident macrophage). We incorporate these two mechanisms in the model as a single growth term. The growth process is upregulated by the presence of activated macrophages (*M*_1_ and *M*_2_), and is independent of the activation state. Macrophage cell loss occurs via natural cell turnover from all states.

##### Activation of macrophages

Activated macrophages can be in an *M*_1_ or an *M*_2_ state. We assume this occurs through both activation from a resting state and re-polarisation between the two activated states. We also assume that the cells activate to the pro-repair *M*_2_ state at a constant rate in early stage disease, for tissue repair and maintenance, and activate to the inflammatory *M*_1_ state when they detect the presence of the endometrial cells [30]. *M*_1_ activation is upregulated by activated NK cells, via the cytokine IFN-γ [14, 22], as detailed below.

##### Activated macrophage actions

M1-type macrophages produce pro-inflammatory cytokines, such as interleukins IL-12 and IL-1 [27]. These are known to activate and stimulate production of NK cytokines. We therefore assume *M*_1_ cells upregulate activation of NK cells. We model M1-type cells as anti-lesion cells that assist in clearance of ectopic endometrial cells, and M2-type cells as pro-lesion cells that promote the growth of lesions once endometrial cells have attached to the peritoneal tissue [16].

#### 2.1.2 Natural killer cells

NK cells exist in one of two states: resting NK cells, denoted by *K*_0_, and activated NK cells, denoted by *K*_*A*_. We model both NK dim and bright cells simply as activated NK cells with moderate cytokine production and cytotoxicity.

##### Resting NK cells

Resting NK cell turnover is through a balance of steady recruitment from the blood stream and loss through natural cell turnover and activation. We assume *K*_0_ cells have insignificant cytokine production and cytotixicity compared to their activated form, *K*_*A*_.

##### Activation of NK cells

NK cell activation occurs through pro-inflammatory cytokines, such as interleukin IL-12, and interferons IFN-α and IFN-γ [14, 49, 50]. We assume the main producers of these are macrophages, particularly IL-12 which is produced at high levels by M1-type macrophages [27]. Consequently we assume that *M*_1_ cells upregulate activation of *K*_*A*_ cells.

##### Activated NK cell action

Activated NK cells are assumed to be responsible for cytokine release, cell clearance, and NK proliferation. This is modelled as upregulation of M1-type polarisation (via IFN-α [14, 22]), clearance of endometrial cells, and proliferation of activated NK (via IL-2 [49]).

##### Activated NK cell clearance

Activated NK cells have a natural turnover rate similar to resting NK cells. Additionally, activated NK cells are known to become exhausted from the execution of cell clearance [51], after which they no longer contribute to cell clearance. To accommodate this, we model *K*_*A*_ to have a higher cell loss rate than *K*_0_ upon exposure to endometrial cells.

#### 2.1.3 Endometrial cells

We define the endometrial cells to be in three states: eutopic (within the uterus, *E*_0_), in (peritoneal) fluid (*E*_*F*_), and attached (to the peritoneum and/or external surface of the reproductive system/gut, *E*_*A*_). *E*_*A*_ cells are early lesion cells which then progress to endometriosis. Transitions between the three cell states are strictly sequential (i.e. an attached endometrial cell will never detach).

##### Eutopic endometrial cells

these are endometrial stromal and epithelial cells located on the internal surface of the uterine cavity. The functionalis layer of the endometrium sheds during menstruation, with stromal and epithelial cells found in menstrual fluid. These cells are cleared from the uterine cavity via ‘normal’ menstruation (cleared via cervix) or enter the peritoneal cavity via retrograde menstruation, where they become fluid cells, *E*_*F*_.

##### Fluid and attached cell turnover

when cells enter the peritoneal cavity via retrograde menstruation, they are first collected within the peritoneal fluid, and hence we denote them to be in the ‘fluid’ cell state. The fluid cells attach to the peritoneum, from which they can form a lesion. Both fluid and attached cells are cleared via natural cell turnover and clearance initiated by the immune system. We assume the same immune clearance rate for fluid and attached cells. Attached cells proliferate to form lesions, and this proliferation is upregulated by the presence of *M*_2_ cells.

##### Endometrial cell action

Fluid and attached endometrial cells are detected by macrophages, and their presence results in activation of resting macrophages to an *M*_1_ state.

### 2.2 Model Equations

Using the assumptions described above, we construct a system of ordinary differential equations. Upregulation terms follow either mass-action or Hill kinetics, and proliferation and production terms are modelled as logistic growth. The cyclic function for eutopic (*E*_0_) endometrial cell influx, *µ*_*E*_(*t*) (Eq. (9)), was fit to normalised menstrual cycle volume data [52]. Below, we present the system of equations, and describe the terms that govern cell dynamics and interactions. The cell dynamics under the default parameter values are shown in Fig. 1(b). Parameter values are given in Supplementary S1.

#### 2.2.1 Macrophage cell dynamics

Equations (1) to (3) describe resting macrophage (*M*_0_), M1-type macrophages (*M*_1_), and M2-type macrophages (*M*_2_) dynamics, respectively.

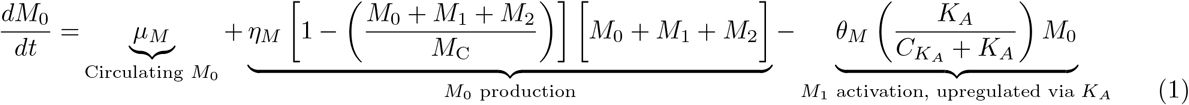

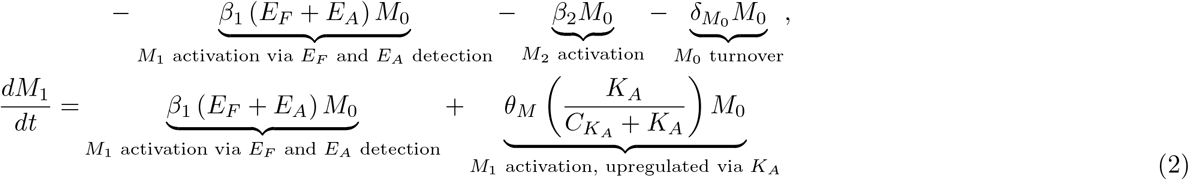

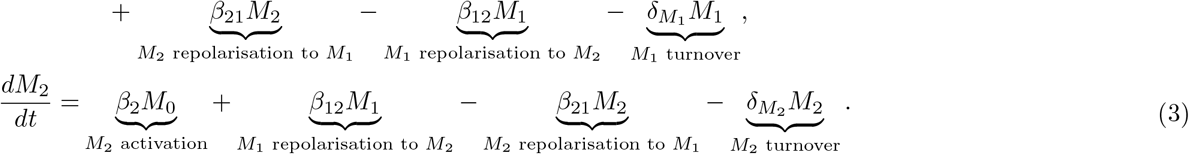

Here, *µ*_*M*_ is the supply rate of *M*_0_ from circulation; 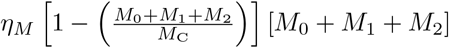 describes *M*_0_ production, with carrying capacity *M*_C_ and rate 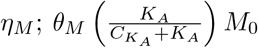 describes *M*_0_ activation to *M*_1_, upregulated by *K*_*A*_, with rate *θ*_*M*_; *β*_1_ (*E*_*F*_ + *E*_*A*_) *M*_0_ describes *M*_0_ activation to *M*_1_, upon detection of *E*_*F*_ and *E*_*A*_, with rate *β*_1_; *β*_2_*M*_0_ describes *M*_0_ activation to *M*_2_ with rate *β*_2_; *β*_21_*M*_2_ describes repolarisation of *M*_2_ to *M*_1_, and *β*_12_*M*_1_ repolarisation of *M*_2_ to *M*_1_; 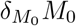 describes the natural turnover of *M*_0_, and similarly for 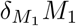 and 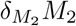.

#### 2.2.2 Natural Killer cell dynamics

Equations (4) and (5) describe resting (*K*_0_), and activated (*K*_*A*_) NK cell dynamics, respectively.

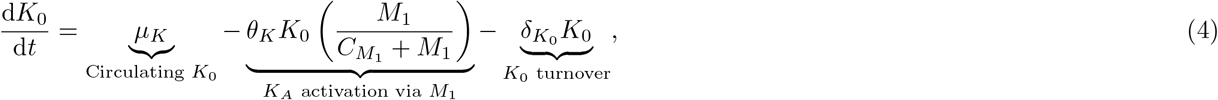

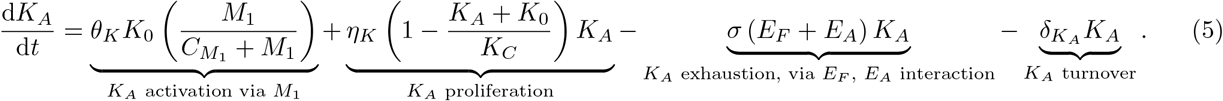

Here, *µ*_*K*_*K*_0_ is the supply rate of *K*_0_ from circulation; 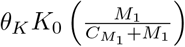 describes *K*_0_ activation to *K*_*A*_, upregulated by *M*_1_, with rate 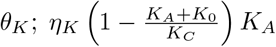 describes *K*_*A*_ proliferation, with carrying capacity *K*_*C*_ and rate *η*_*K*_; *σ* (*E*_*F*_ + *E*_*A*_) *K*_*A*_ describes *K*_*A*_ exhaustion, due to *E*_*F*_ and *E*_*A*_ clearance, with rate *σ*; 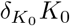 and 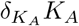 describe the natural turnover of *K*_0_ and *K*_*A*_ respectively.

#### 2.2.3 Endometrial cell dynamics

Equations (6) to (7) describe eutopic (*E*_0_), in peritoneal fluid (*E*_*F*_), and attached (*E*_*A*_) endometrial cell dynamics respectively.

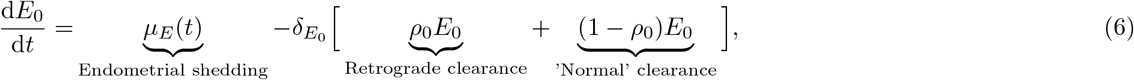

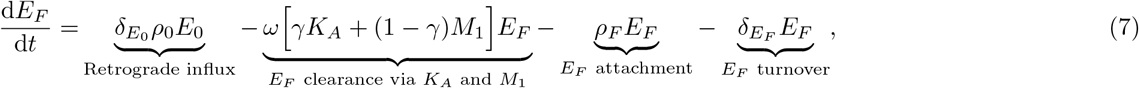

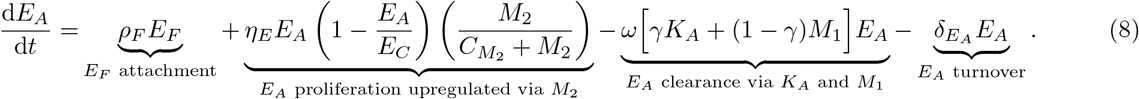

Where:

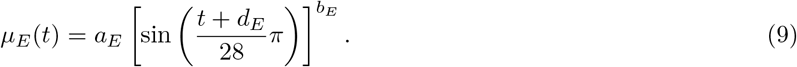

Here, *µ*_*E*_(*t*) describes endometrial shedding; 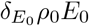 is *E*_0_ retrograde clearance to *E*_*F*_, and 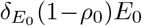 is ‘normal’ *E*_0_ clearance; *ρ*_*F*_ *E*_*F*_ represents *E*_*F*_ attachment, becoming *E*_*A*_, with rate *ρ*_*F*_; *ω* [*γK*_*A*_ + (1 − *γ*) *M*_1_] *E*_*i*_ describes *E*_*F*_ (*i* = *F*) and *E*_*A*_ (*i* = *A*) clearance, via *K*_*A*_ and *M*_1_, with rate *ω* and *γ* the proportion of *K*_*A*_ relative to *M*_1_ cell clearance; 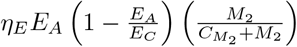 describes *E*_*A*_ proliferation, with carrying capacity *E*_*C*_, upregulated by *M*_2_, with rate 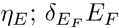 and 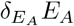 describe the natural turnover of *E*_*F*_ and *E*_*A*_, respectively.

#### 2.2.4 Constant influx surrogate model

Taking 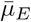 as the average influx over the cycle, we define a surrogate model for the system in which we assume a constant influx into *E*_0_, i.e.:

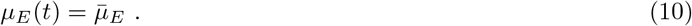

This results in a sufficient match between the constant influx and the cyclic influx systems, in terms of both average quantities and emergent behaviour. The surrogate model provides two advantages to the cyclic influx model: 1) it is computationally more efficient, and 2) it allows for clearer analysis of some aspects of the emergent system dynamics.

#### 2.2.5 Parameters and initial conditions

Unless otherwise specified, the parameters for the model are given in Supplementary S1, and the response of the system for these parameter values is shown in Fig. 1(b). Where possible, we use parameters from previous within-host immune dynamics models [19, 20, 53], or estimate them using *in-vitro, in-vivo*, or clinical data [5–7, 15, 17, 25, 52, 54–58]. Due to a lack of quantitative data on immune cell-endometrial cell interactions, several parameters were taken from the cancer and viral literature. Details on the parameterisation are given in Supplementary S2.

To solve the initial value problem for a particular parameter set, we first solve the system of equations using the surrogate model (Section 2.2.4), whose steady state solutions inform the initial conditions for the full system with cyclic influx. The initial conditions for the surrogate model solution use a combination of model steady state values and observations of cell concentrations in the literature [5, 25, 56] (see Supplementary S1).

### 2.3 Characterisation of disease dynamics

#### 2.3.1 Attachment and disease definitions

We use the value of *E*_*A*_ at the end of an endometrial cycle (28 day cycle) to determine if the response results in a diseased state. A low/no disease system is defined as end of cycle attachment below some threshold value, *ε*: *E*_*A*_(*t* = 28*k*) *< ε, k* ∈ ℤ_≥0_, and *t > τ*, where *τ* is the initial transient time for the system to reach a cyclic state, for some *ε* ∈ ℝ_≥0_.

In addition to classifying responses as high disease or low/no disease, we consider different response states across the endometrial cycle. We classify system dynamics into three attachment states based on the threshold value *ε*:

1. Low attachment (low/no disease): cell attachment is always below the threshold, *E*_*A*_(*t*) ≤ *ε*∀ *t > τ*;
2. Transient attachment (low/no disease): peak cell attachment is above the threshold but decreases to below the threshold at the end of the cycle, *Ê*_*A*_ *> ε, E*_*A*_(*t* = 28*k*) ≤ *ε* ∀*k* ∈ ℤ, *t > τ*;
3. Sustained attachment (diseased): end of cycle attachment is above the threshold, *E*_*A*_(*t* = 28*k*) *> ε* ∀*t > τ*; where *Ê*_*A*_ is the maximum *E*_*A*_ value over the menstrual cycle, and *E*_*A*_(*t* = 28*k*) the number of attached cells at the end of the endometrial cycle after initial transient time *τ* days. The term low/no disease is used as the nature of the modelling paradigm does not permit *E*_*A*_ = 0, for *t >* 0 days. Examples of each of these attachment states are given in Fig. 2.

**Figure 2:**
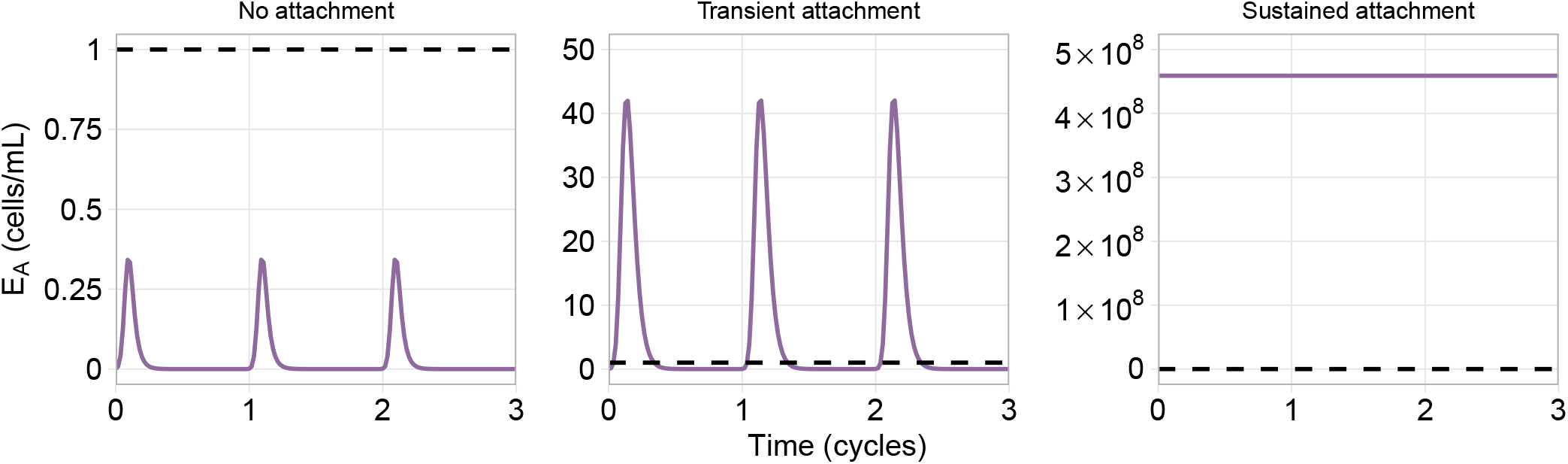
Examples of the three emergent attachment state dynamics observed, assuming a threshold of *ϵ* = 1 cell/mL (indicated by the black dashed line in the plots). The left two plots (no attachment and transient) are hypothesised to be the low/no disease states, and the right plot (sustained) is hypothesised to be the diseased state.

#### 2.3.2 Parameters and response metrics of interest

To investigate the questions of interest around endometriosis lesion onset, we examine the model’s response to the variation of three parameters: the proportion of eutopic endometrial cells that are shed retrograde into the peritoneal fluid (*ρ*_0_), macrophage detection rate of endometrial cells (*β*), and the clearance rate of the endometrial cells by the immune cells (*ω*). In our model, macrophages’ ability to detect endometrial cells represents the level of apoptotic marker expression and viability of the endometrial cells. M1-type macrophage activation, which occurs predominantly through endometrial cell detection (*β*), and the subsequent activation of NK cells is a pre-requisite to cell clearance (*ω*). The primary system responses we are interested in are: the number of attached endometrial cells (*E*_*A*_) at the end of the cycle, as an indicator for the level of disease; and the level of macrophage activation, measured as the proportion of total macrophages in each activation state *M*_1_ and *M*_2_, denoted by 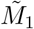 and 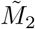 respectively, as a comparison to immune profiles observed in endometriosis patients. We use proportions for macrophage activation to allow for simple comparison with studies on peritoneal fluid cell compositions, which report the proportions of macrophages expressing specific markers.

### 2.4 Code availability

The code to generate these results is openly available at https://github.com/clairemiller/math_modelling_immune_endo. The ODE system was simulated Matlab’s ode15s (R2024b) [59], and the bifurcation analysis was performed using the software AUTO-07p [60].

## 3 Results

### 3.1 Low attachment analysis

We perform an eigenvalue analysis to identify the stability of each of the steady states. By requiring the disease-free steady state to be stable, we find that the following inequality must be satisfied:

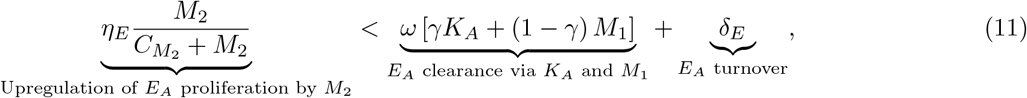

where *η*_*E*_ is the growth rate of endometrial cells, *C*_*M*_2 is the action limiting capacity for *M*_2_ upregulation of *E*_*A*_ proliferation, *ω* is the lysis rate of *E*_*A*_ by the immune cells, with *γ* representing the proportion of *K*_*A*_ relative to *M*_1_ lysis, and *δ*_*E*_ is the natural clearance rate of *E*_*A*_ (see Supplementary S1 for parameter values). The full analysis is given in Supplementary S4. The biological interpretation of this disease-free state is that the proliferation rate of the attached endometrial cells, upregulated by *M*_2_, must be smaller than the clearance rate of endometrial cells—both cell clearance due to the immune response and natural turnover.

This analysis only provides information for when the system is in the disease-free state, and does not provide insight into how a transition to disease may occur.

### 3.2 Hypotheses of disease onset

#### 3.2.1 Retrograde influx

We are interested in understanding whether the observed immune profile of peritoneal fluid from endometriosis patients could emerge due to the presence of the endometrial cells with no immune dysfunction, and if increased endometrial presence is associated with disease (Hypothesis A). To investigate this, we explore the response of the system as the level of retrograde influx into the peritoneal fluid increases. This is done by increasing *ρ*_0_, the proportion of menstrual debris that clears retrograde from the uterus.

The system response to variations in the retrograde influx is plotted in Fig. 3. This figure shows the dynamics of the attached endometrial cells and activated immune cells. High levels of retrograde influx is associated with an increase in the proportion of macrophages activated to M1-type (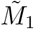, Fig. 3(c)) and the concentration of activated NK cells (*K*_*A*_, Fig. 3(a)), but there is no significant change to the proportion of macrophage activated to M2-type (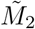, Fig. 3(d)). In our model, *M*_2_ activation is only indirectly affected by endometrial cell presence, as a result of the subsequent changes in *M*_0_ and *M*_1_ levels, which are not large enough here to significantly affect 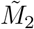. The increase in 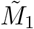, and consequently *K*_*A*_, is attributed to the upregulation of *M*_1_ in response to the presence of *E*_*F*_ and *E*_*A*_, as described in Supplementary S1. We also observe that the relaxation time of *M*_1_ and *K*_*A*_ is longer than the menstrual cycle length, resulting in a sustained state of inflammation. This result aligns with the description of endometriosis as a chronic inflammatory disease [2, 22].o

**Figure 3:**
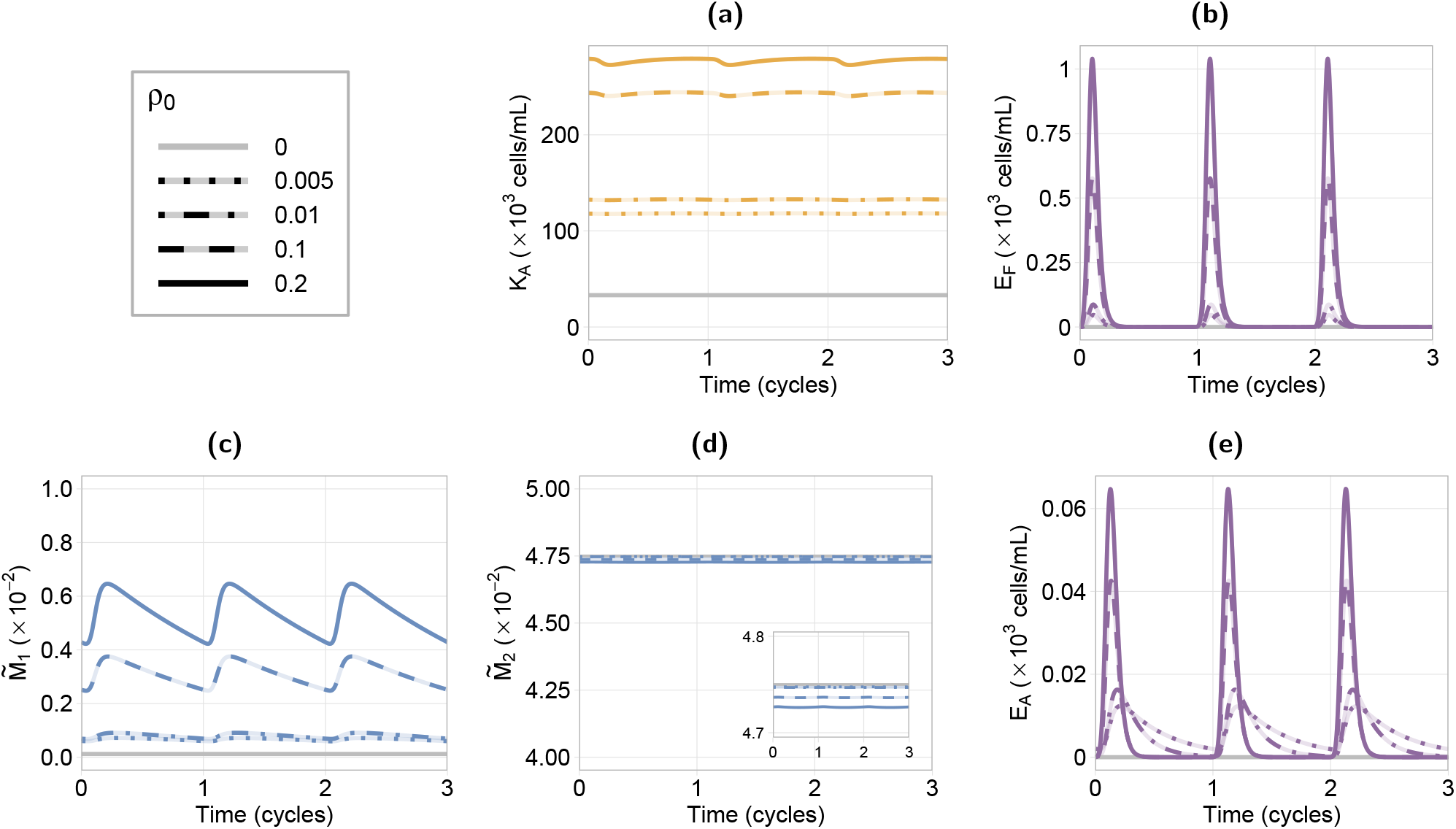
Timeseries of the response for varying endometrial retrograde influx (*ρ*_0_) shown for three menstrual cycles. We see large peaks in the endometrial cells, both in fluid (*E*_*F*_, (b)) and attached (*E*_*A*_, (e)) with the menstrual phase of the cycle. Oscillations are also seen in the proportion of macrophage activated to *M*_1_ type, (c), and NK cell activation, (a). Increased influx causes an increase in the proportion of macrophage activated to *M*_1_ and activated NK cells, but does not have a significant effect on the proportion of macrophages activated to *M*_2_, (d). Results show high retrograde influx is associated with a small sustained increase in inflammation but not sustained attachment.

We see that a higher retrograde influx results in an increase in the peak level of endometrial cell attachment Fig. 3(e). Contrary to expectations, low influx leads to higher attachment levels at the end of the cycle (see *ρ*_0_ = 0.005, dotted line in Fig. 3(e)). This is because an increased influx of endometrial cells causes an increased activation of *M*_1_ cells, and consequently *K*_*A*_ cells, as noted above. As a result, immune clearance of the endometrial cells is increased. At low levels of *ρ*_0_, there is a decrease in the activation of *M*_1_ cells, and consequently a decrease in the immune clearance of endometrial cells. The proliferation rate of the attached endometrial cells is comparable between the different influx levels, as increasing *ρ*_0_ has an insignificant effect on the *M*_2_ cell levels. Consequently, the low influx systems maintain a low level of sustained attachment while high influx systems exhibit only transient attachment.

Therefore, we see that our results show an increase in retrograde influx is associated with a small, sustained increase in inflammation, but is not associated with increased disease, opposing Hypothesis A.

#### 3.2.2 Immune dysfunction

There are several hypotheses regarding immune system dysfunctions in endometriosis pathogenesis. Two immune dysfunctions we explore are altered detection (Hypothesis B1, model parameter *β*_1_) and clearance (Hypothesis B2, model parameter *ω*) of the endometrial cells. We consider a reduction in clearance rate by both activated immune cell types, *M*_1_ and *K*_*A*_, noting that NK cells are assumed to be the main clearance cell in the model. Detection dysfunction in our model only applies to macrophages as we assume they are the predominant cell type responsible for detecting the endometrial cells. Importantly, detection dysfunction also has a subsequent effect on cell clearance, through a reduction in immune cell pro-inflammatory activation, which is a prerequisite for cell clearance.

Five example responses for moderate retrograde influx are shown in Fig. 4(a). Examples 1 and 5 in Fig. 4(a) show a healthy and diseased response respectively. In the healthy response, both *E*_*A*_ and *E*_*F*_ are cleared by the end of the cycle, while in the diseased response, only *E*_*F*_ is cleared, and *E*_*A*_ has a high level of sustained attachment.

**Figure 4:**
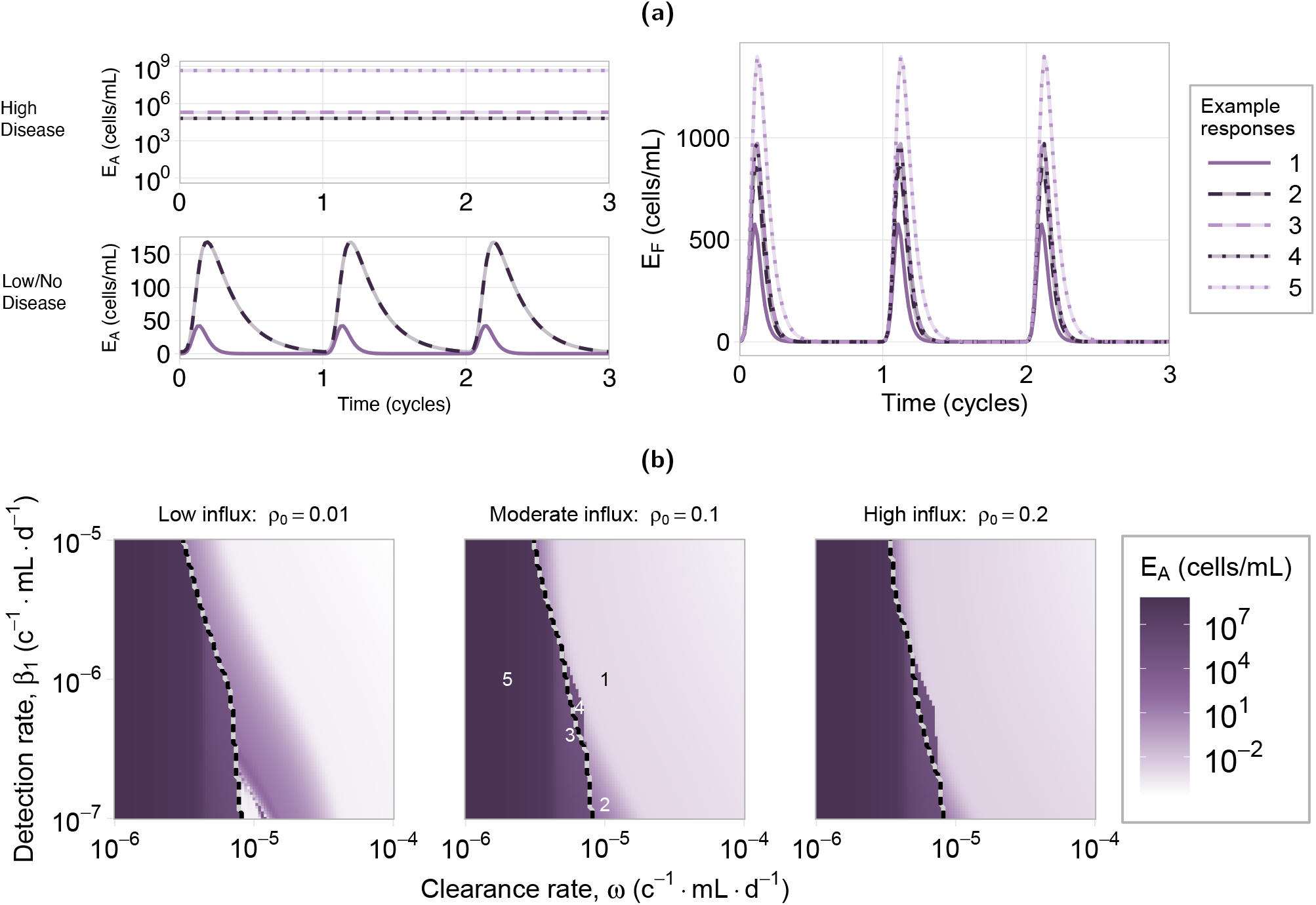
Endometrial attachment and in fluid response to immune dysfunction. (a) Example responses of attached and fluid endometrial cell dynamics at locations indicated on the heatmaps in (b) (moderate influx). (a) top left: the *E*_*A*_ response for the high attachment (high disease) examples. (a) bottom left: the *E*_*A*_ response for the low attachment (low/no disease) examples. (a) right: the level of endometrial cells in fluid, *E*_*F*_, which we observe are always cleared by the end of each cycle. (b) Heatmap showing the level of attached endometrial cells at the end of a cycle. The dashed line indicates the partition between high and low attachment regions according to the inequality in Eq. (11). Units (c^−1^ · mL · d^−1^) are short for (cells^−1^· mL · day^−1^). These results indicate that reduced immune clearance ability is a stronger driver of disease compared to reduced detection ability.

The endometrial cell response to variations in both immune detection and clearance ability is shown in Fig. 4(b) for three levels of retrograde influx, *ρ*_0_ (low, moderate, and high). A similar investigation was also performed for different levels of endometrial cell attachment rate, which showed attachment rate does not affect these results (see Supplementary S4). Figure 4(b) shows the end of cycle (i.e. cycle day 28) values for endometrial attachment (*E*_*A*_). We see that, for all *ρ*_0_ levels, *E*_*A*_ transitions from a low attachment state (low/no disease, light purple) to a high attachment state (high disease, dark purple) with decreasing clearance ability. Over a wide range of detection rates, *β*_1_, a reduction in clearance rate, *ω*, will result in the disease transition. However, for reductions in the detection rate, *β*_1_, this transition only occurs across a narrow range of *ω* values. This shows the disease transition is dominated by the immune clearance rate, *ω*.

The dashed lines in Fig. 4(b) (and Fig. 5(a), Fig. 5(b)) partition the regions governed by the inequality in Eq. (11). This partition is calculated using the surrogate model (see Section 2.2.4). Responses in the right region are predicted to exhibit low attachment behaviour, and those to the left, high attachment behaviour. The alignment between this partition and the disease transition in the numerical results indicates that this relationship is a suitable indicator of disease. The region of disagreement showing high *E*_*A*_ but predicted to be in low-disease region is a region where attached endometrial cell proliferation and removal are approximately equal (see example responses 3 and 4 in Fig. 4(a)).

**Figure 5:**
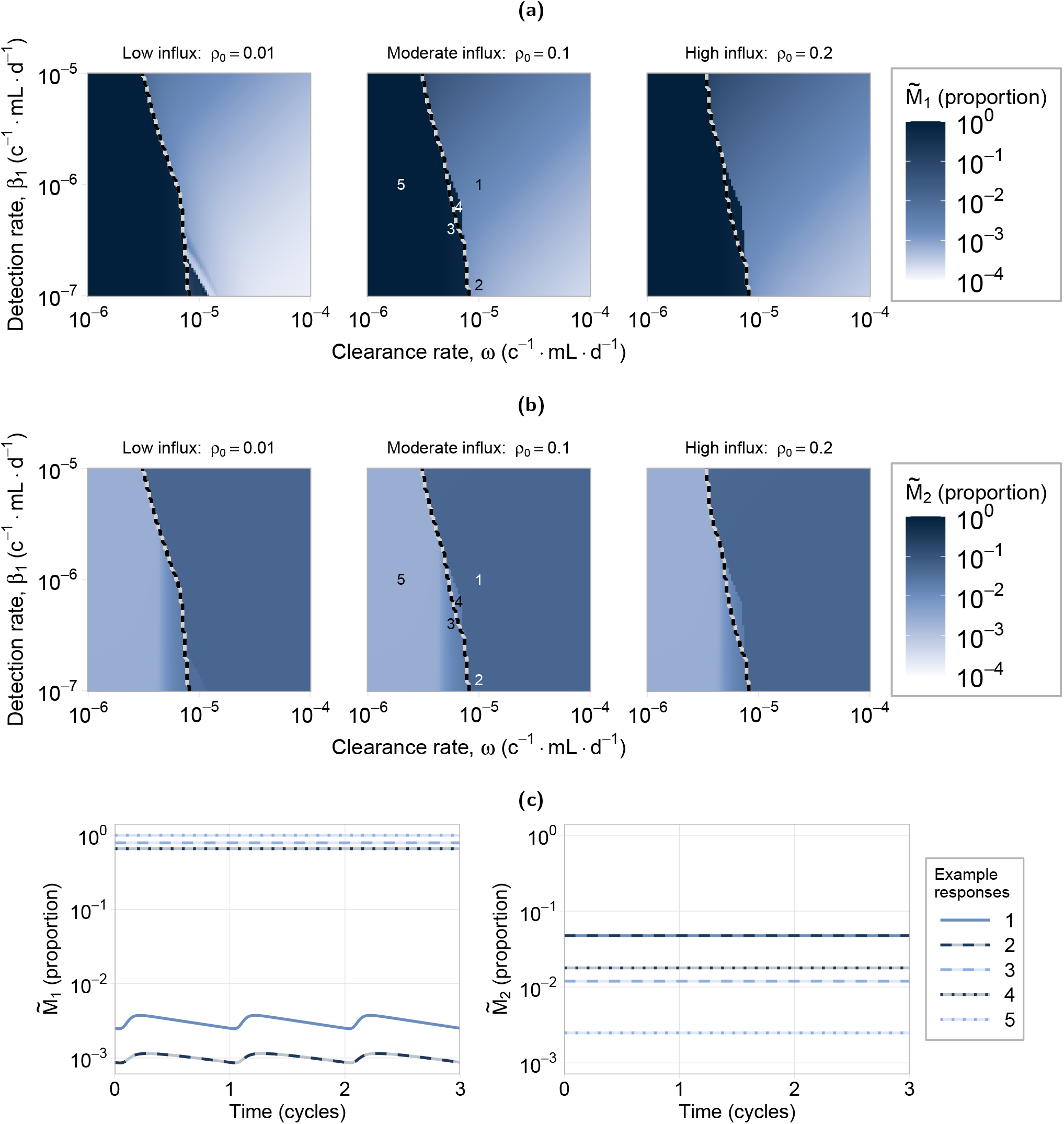
Macrophage activation response to immune dysfunction. (a), (b) Heatmaps showing the proportion of macrophage polarised to *M*_1_ and *M*_2_, respectively, at the end of a cycle. The dashed line indicates the partition between high and low attachment regions according to the inequality in Eq. (11). Units (c^−1^ · mL · d^−1^) are short for (cells^−1^ · mL · day^−1^). (c) Example responses at locations indicated on the heatmaps in (a), (b). (c) left: the proportion of macrophages in an *M*_1_ activation state. (c) right: the proportion of macrophages in an *M*_2_ activation state. Note examples 1 and 2 are overlapping in right plot. The high attachment region is associated with high *M*_1_ activation and low *M*_2_ activation. In the low attachment region, decreasing clearance rate is associated with an increase in *M*_1_ activation while decreasing detection rate is associated with a decrease in *M*_1_ activation. There is no significant effect on *M*_2_ activation in the low attachment region.

Figures 5(a) and 5(b) show the macrophage activation response to a reduction in immune clearance and detection rate. In both figures, in the high attachment region, there is a high proportion of *M*_1_ activation and a reduced proportion of *M*_2_ activation as a result of the high endometrial cell presence. This transition can be seen in the Fig. 5(c), with low 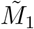 and high 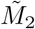 in the two low attachment example responses 1 and 2, and high 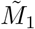 and low 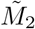 for the high attachment example responses 3–5.

In the low attachment region, the results show an increase in 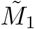 with decreasing clearance ability, while decreasing detection ability results in a decrease in 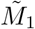. The latter can be seen in example response 2 in Fig. 5(c), which shows a decrease in 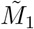 but no change to 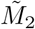 compared to example response 1. A decreased detection ability reduces the rate at which macrophages activate into an *M*_1_ state, consequently leading to this reduction in inflammation.

These results show that a reduced immune clearance rate, *ω*, leads to high disease and high *M*_1_ activation. Neither hypothesis alone leads to an increase in *M*_2_ activation in our results. However, elevated *M*_2_ activation may occur when *M*_2_ is upregulated by *E*_*A*_, but only once the system already exhibits a high disease state (see Supplementary S6). Consequently, our results show the reduced clearance hypothesis, Hypothesis B2, is most consistent with the observations of increased macrophage activation in endometriosis patients compared to both the reduced detection and increased retrograde hypotheses.

### 3.3 Disease transition

We now consider the case, for Hypothesis B, where the immune clearance and detection have a gradual decline in function, rather than a pre-existing dysfunction. This can be explored through a bifurcation analysis in *ω* and *β*_1_. A bifurcation analysis can be used to determine the equilibrium of a system as model parameters change, and identify points at which the behaviour of these equilibrium changes between being stable, bistable, and unstable, for example. This allows us to identify regions in parameter space that can sustain both a high disease and a low/no disease state (bistability). The observed transition from low to high endometrial attachment in Fig. 4(b) with decreased immune function is indicative of bistable system dynamics.

We perform a bifurcation analysis, using the surrogate model described in Section 2.2.4. Figure 6(a) shows the biologically observable states, from this analysis, of the attached endometrial cells for varying clearance (left) and detection (right) rate. We reclassify solutions containing limit cycles (using the median value of 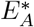, shown as a dashed line in Fig. 6(a), where 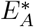 is a solution containing a limit cycle) as their period of oscillation exceeds the expected timescale of disease onset in the presence of the innate immune response only, beyond which the adaptive immune response is known to play a role [2]. Complete bifurcation diagrams can be found in Supplementary S5.

**Figure 6:**
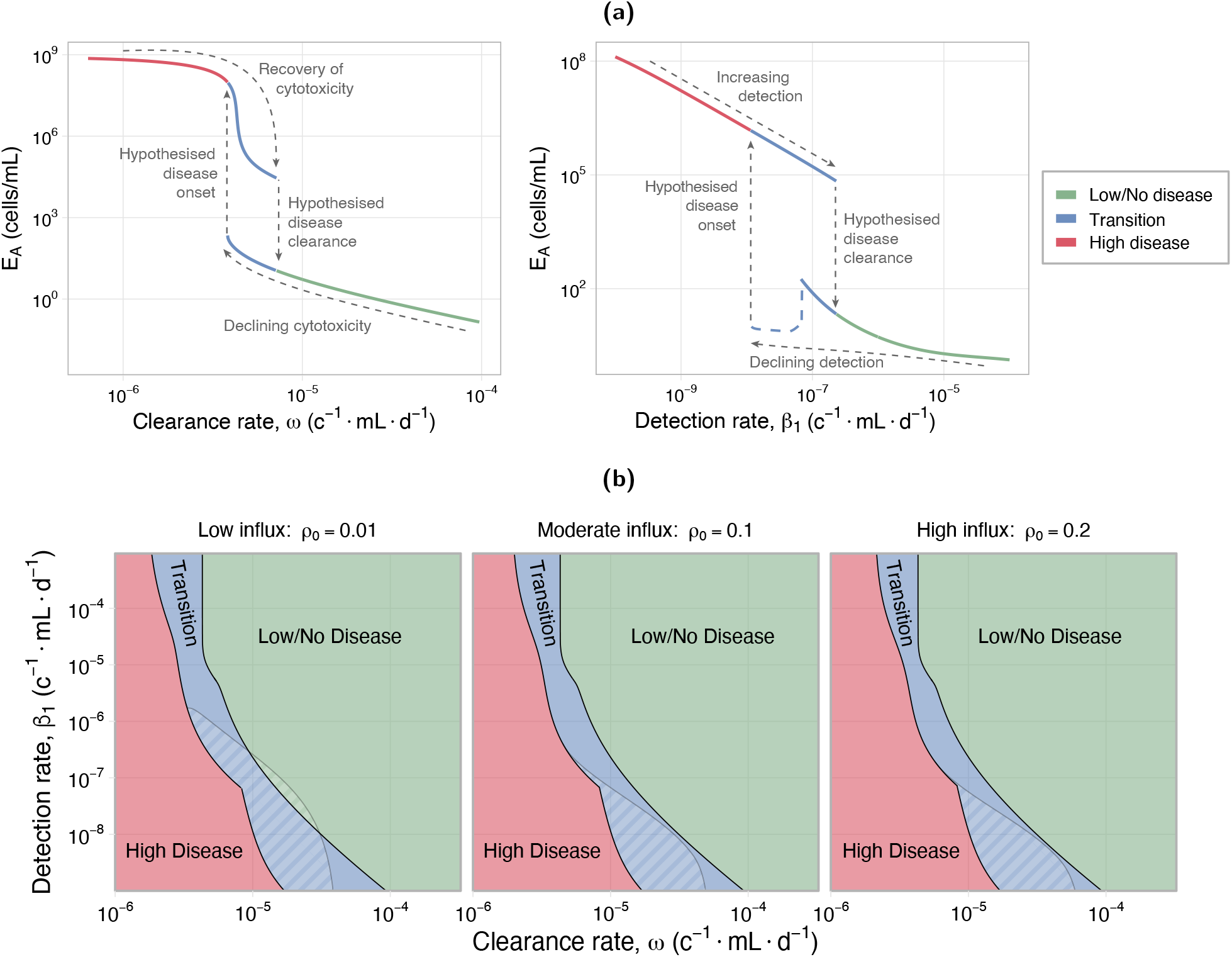
Disease transition plots for clearance and detection rate: (a) clearance (left) and detection (right) rate independently. The dashed blue line (right plot) indicates the solutions reclassified from a limit cycle—showing the median value of 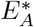. (b) Co-dimensional bifurcation in clearance (*ω*) and detection (*β*_1_) rate. Hatched region shows the reclassified limit cycle solutions. The complete bifurcation diagrams are given in Supplementary S5. Fixed values for (a) are *β*_1_ = 10^−6^ cells^−1^ ·mL· day^−1^, *ω* = 10^−5^ cells^−1^ ·mL ·day^−1^, and *ρ*_0_ = 0.1. Units (c^−1^ ·mL ·d^−1^) are short for (cells^−1^ ·mL ·day^−1^). A patient can start in the disease-free region (green), and then transition through the bistable region (blue), still in low/no disease, but then switch into the high disease state either due to a decline in immune function, or due to additional insult to the system. Disease clearance from this state requires significant improvement to immune function.

For both clearance and detection rate, we observe a bistable regime where the system exhibits hysteresis, which we term the transition region, shown in blue on Fig. 6(a). Biologically, this indicates that a healthy individual can start in a low/no disease state (green); upon an initial decline in immune function, they enter the transition region, remaining in the low disease state (bottom blue); with further immune function decline, or other changes to the endometrial or immune cell states, the individual will switch into a high disease state (red, or top blue, respectively), which we hypothesis to be disease onset. The individual is unable to recover from this diseased state with subsequent improvement in immune function (top blue), unless the improvement achieves the hypothesised disease clearance indicated on the figure.

We perform a co-dimensional bifurcation analysis, using the surrogate model, to look at both clearance and detection rate simultaneously. A full co-dimensional bifurcation diagram is provided in Supplementary S5.3.The biologically observable states from this analysis are shown in Fig. 6(b) for three levels of retrograde influx (low, moderate, and high), with solutions containing limit cycles reclassified as described above (hatched region in Fig. 6(b)). Regardless of the level of retrograde influx, we observe similar behaviours – changing the influx level simply shifts the transition region in parameter space. This reinforces our finding that increased retrograde influx is neither associated with increased disease, nor a driving factor of system dynamics in our model. Notably, a transition region is always present between the low/no and high disease states. As detailed previously, this indicates that a switch into high disease (disease onset) is not recoverable without significant improvements in immune system function (disease clearance).

## 4 Discussion

In this study we develop a novel compartmental mathematical model of the innate immune response to endometrial cell influx from retrograde menstruation, to describe the early stages of superficial peritoneal endometriosis lesion onset. We use our model to interrogate two key questions around endometriosis onset. Firstly, could increased endometrial cell presence describe altered immune states observed in endometriosis patients and lead to disease (Hypothesis A). Secondly, of two hypothesised disorders: immune system detection and clearance of endometrial cells in menstrual debris (Hypotheses B1 and B2), which is more associated with disease and consistent with clinical observations. This is the first mathematical model that has been developed to interrogate mechanisms of endometriosis lesion onset, highlighting the importance of the immune response.

Our results show that an increase in retrograde menstruation, with no immune system dysfunction, is not associated with increased disease, but is associated with a small, sustained increase in inflammation (Fig. 3). We find the key driver of disease onset is a decrease in the clearance rate of endometrial cells by immune cells (Fig. 4), supporting Hypothesis B2. Both decreased clearance rate and high disease are associated with an increase in M1-type macrophage activation, but not M2-type activation (Fig. 5). This increase is sustained across the cycle and may cause a state of chronic inflammation. A decrease in immune detection rate, which represents reduced phagocytosis and apoptotic marker expression by the endometrial cells, will also result in disease onset within a specific range of clearance rates, indicating that different immune profiles can be indicative of disease. Given an immune profile, we provide a relationship between attached endometrial cell proliferation and removal that is a predictor of disease (Eq. (11)). Through a bifurcation analysis, we show that a transition region exists that dictates the dynamics of disease onset and disease clearance (Fig. 6). This result describes how, following disease onset, a significant recovery of immune function is required to obtain disease clearance. Our analysis supports a hypothesis that a decline in immune function, in particular immune cell cytotoxicity, such as due to immune exhaustion as a result of chronic inflammation [61], may be a driver of disease onset. We note that our results show a sustained increase in inflammation (*M*_1_), in the low/no disease state, under both the reduced clearance hypothesis and the increased retrograde influx hypothesis.

A recent review has challenged the hypothesis that frequency and volume of retrograde menstruation is different between endometriosis patients and controls [8]. This is supported by our results, which show no association with increased retrograde influx and disease (Fig. 3). Endometrial stem and progenitor cell concentrations in peritoneal fluid have been observed to be one to two orders of magnitude higher during the menstrual phase, compared to non-menstrual phase, in both control and endometriosis patients [13]. Endometriosis cases had increased variation between patients, with some patients measuring concentrations comparable to control cases, and others measuring significantly higher cell concentrations, which may indicate there is no unique pathway to disease onset. This observation does not support a particular hypothesis based on our results: a significant increase in endometrial cells in fluid can be seen during the menstrual phase for all levels of retrograde influx (Fig. 3), and our bifurcation analysis results show only small changes in endometrial cells in fluid between low/no disease and high disease, particularly in the transition region (Supplementary S5). This example demonstrates that, without high precision on cycle time and additional physiological information, single time point data on peritoneal fluid cell concentrations is insufficient to inform mathematical models or indicate the presence of disease.

Several studies have shown increases in both M1- and M2-type macrophages, as proportions of total macrophage, in endometriosis patients [16, 17, 62]. Our results show an increase in M1-type macrophages is associated with increased retrograde menstruation (Fig. 3(c)) and decreased immune clearance capabilities (Fig. 5(c)); and a decrease in M2-type macrophages is associated with a decrease in immune clearance capabilities (Fig. 5(c)). Our model does not predict an increase in both macrophage activation states under any hypothesis. The lack of increased M2-type activation in our results is due to no *M*_2_ upregulation mechanism in the model. By incorporating *M*_2_ upregulation by *E*_*A*_, a high-disease state, characterised by both increased *M*_1_ and *M*_2_ activation, may occur (see Supplementary S6). However, this upregulation alone is not sufficient to trigger disease onset. It is important to also note that clinical data is only collected after confirmed diagnosis, which is usually several years after disease onset [1, 63]. This leads to ambiguity in which observations of increased M1- or M2-type activations are associated with the early stages of disease and those that emerge during later stages of lesion development and maintenance.

Few studies have focused on NK cell concentrations and activation types in peritoneal fluid. A significant increase (approximately four-fold) in the proportion of leukocytes that stained for NHK-1 in Stage I disease has been observed [6], which could indicate an increase in the concentration of activated NK cells. Such an increase is predicted by our model under the increased retrograde menstruation hypothesis, and the decreased cell clearance hypothesis in the low disease state (see Supplementary S5).

To our knowledge, this is the first mechanistic mathematical model developed in the context of endometriosis. Our model combines understanding of macrophage, NK cell, and endometrial cell dynamics. Our model provides a framework that can be used to provide insight into open questions, such as those considered in this paper, around system changes that contribute to disease onset compared to those that emerge due to the disease. It can also be used to better understand the interacting role of different mechanisms hypothesised through *in vitro* models within an *in vivo* environment.

There are several known modelling limitations to this work. We use a deterministic model to describe interactions and transitions, meaning that stochastic effects, such as menstrual cycle variation, low cell numbers, and delays in immune onset, are not accounted for. The cell transitions and interactions are based on understanding of immune interactions from the endometriosis, cancer, and viral literature, as there is limited understanding of interactions between immune cells and endometrial cells. As with many models, there is also little quantitative knowledge of the system, particularly for the early stage of lesion onset. This is in part due to the extensive delays patients experience before they receive a clinical diagnosis for endometriosis [63]. Our model has highlighted the need for several quantitative data requirements. In particular, longitudinal studies of patients that capture time-series immune profiles, and connected data sets that provide full peritoneal fluid immune cell profiles of individuals, and also record menstrual cycle day at time of collection. In the future we will explore opportunities around parameter estimation using clinical and experimental data. For example, our model would greatly benefit from *in vitro* studies of activation states and clearance rates using co-culture assays, animal models, or organoids to measure apoptosis or immune activation markers over time. Future work could also leverage single-cell gene profiling studies [64, 65], alongside an extension of our model that focuses on specific macrophage cell subtypes, to perform parameter estimation.

We have focused our investigation on the innate immune system (though it is known that the adaptive system also plays a role [2]), and dysfunctions related to early-stage disease and the pro-inflammatory immune response. We assume a binary classification of macrophage activation is sufficient to capture the average behaviour of the macrophage activation spectrum: macrophage activation and phenotype dynamics would merit its own modelling investigation prior to inclusion in this model. Additionally, we did not distinguish between the different endometrial cell types, such as stromal fibroblasts, which have been implicated in disease [36, 37]. Further innate immune cells that may also play a role include neutrophils, mast cells, and dendritic cells [2]. Future work will extend the model to include 1) hormonal effects on immune and endometrial cell behaviours, 2) influx of immune cells into the peritoneal fluid as a component of the menstrual debris, 3) investigations into interactions between specific endometrial cell types and the immune cells and 4) explicitly modelling the production and action of particular pro- and anti-inflammatory factors (including cytokines). This will allow us to investigate hypotheses of disease onset relating to progesterone resistance of the endometrial cells [46, 47], hormonal effects on the immune cells [22], abnormal immune function in eutopic endometrium [35, 66], and the effect of altered cytokine production or response on disease onset [3, 14, 21, 31]. With further development and validation, a model such as this could be used for personalised medicine applications. By collecting individual-specific measurements for key parameters it will be possible to develop a risk profile for their susceptibility to disease, or determine the efficacy of potential treatments for preventing lesion onset and growth.

There are many unanswered questions regarding disease onset in endometriosis pathophysiology, including contradicting evidence between different studies and uncertainties about mechanisms that drive disease onset versus those that result from the disease. Our study uses mathematical modelling to provide insight into the role of the innate immune system in superficial peritoneal endometriosis lesion onset. Results show that increased retrograde influx results in small changes to immune profiles but is not associated with disease. Immune dysfunction does lead to disease onset according to our model, in particular a dysfunction in immune clearance ability, and recovery from disease requires significant improvements to immune function. Our model is ideally placed to provide better understanding towards the role of different mechanisms hypothesised through *in vitro* models within an *in vivo* context. With further development and robust parameterisation methods, mathematical modelling will facilitate the identification of potential biomarkers and immunotherapies for endometriosis, and improve our understanding of between-patient variation, moving towards the goal of personalised medicine.

## Acknowledgements

The authors thank Prof. Federico Frascoli for advice on bifurcation analysis and assistance with AUTO-07p. Dr Claire Miller thanks the Aotearoa Foundation, who provided the financial support for the larger project this study is part of, through the Aotearoa Fellowship.

## Notes

https://github.com/clairemiller/ImmuneModelEndometriosis

### Competing Interest Statement

The authors have declared no competing interest.

### Summary of Updates

This version of the manuscript has been revised to include additional supplementary material that show the system response under different endometrial cell attachment rates, and the effect of upregulation of M2-type macrophages by attached ednometrial cells on system dynamics.

