## Supplementary Information S1-6 for "Mathematical modelling of macrophage and natural killer cell immune response during early stages of peritoneal endometriosis lesion onset"

### S1 Parameter tables

| Parameter | Description | Value | Source and type |
|---|---|---|---|
| $\mu_M$ | Constant influx $M_0$ from circulation | $10^4$ cells $\cdot$ mL$^{-1}$ $\cdot$ day$^{-1}$ | See Supplementary Section S2.2 |
| $\mu_K$ | Influx $K_0$ from circulation | $10^4$ cells $\cdot$ mL$^{-1}$ $\cdot$ day$^{-1}$ | See Supplementary Section S2.3 |
| $\eta_E$ | Growth rate of $E_A$ | 1.0 day$^{-1}$ | [54], *In vitro* endometrial cells |
| $\eta_K$ | Growth rate $K_A$ | 0.02 day$^{-1}$ | See Supplementary Section S2.3 |
| $\eta_M$ | Growth rate $M_0$ | 0.2 day$^{-1}$ | [19], Mathematical model (viral) |
| $\delta_M$ | Clearance rate M | 0.02 day$^{-1}$ | [19, 20], Mathematical models (cancer, viral) |
| $\delta_K$ | Clearance rate K | 0.02 day$^{-1}$ | [55], *In vivo* data (macques) |
| $\delta_{E_0}$ | Clearance rate of $E_0$ | 1.0 day$^{-1}$ | [52], Human data |
| $\delta_E$ | Natural clearance rate of E | 0.14 day$^{-1}$ | Guess |
| $\sigma$ | $K_A$ exhaustion due to E | $10^{-5}$ cells$^{-1}$ $\cdot$ mL $\cdot$ day$^{-1}$ | Guess |
| $M_C$ | M carrying capacity | $10^6$ cells/mL | [20], Mathematical model (cancer) |
| $K_C$ | K carrying capacity | $10^6$ cells/mL | Guess |
| $E_C$ | $E_A$ carrying capacity | $10^9$ cells/mL | [20], Mathematical model (cancer) |
| $C_{M_1}$ | $M_1$ action limiting capacity | $10^5$ cells/mL | [53], Mathematical model (cancer) |
| $C_{M_2}$ | $M_2$ action limiting capacity | 100 cells/mL | [53], Mathematical model (cancer) |
| $C_{K_A}$ | $K_A$ action limiting capacity | $10^6$ cells/mL | Guess |
| $\beta_1$ | $M_1$ diff. from $M_0$ due to detection of E | $10^{-6}$ cells$^{-1}$ $\cdot$ mL $\cdot$ day$^{-1}$ | [19, 53], Mathematical models (cancer, viral) |
| $\beta_2$ | $M_2$ diff. from $M_0$ | $10^{-3}$ day$^{-1}$ | [20], Mathematical model (cancer) |
| $\beta_{12}$ | Polarisation from $M_1$ to $M_2$ | $5.0 \times 10^{-5}$ day$^{-1}$ | [19, 20], Mathematical model (viral) |
| $\beta_{21}$ | Polarisation from $M_2$ to $M_1$ | $5.0 \times 10^{-5}$ day$^{-1}$ | [20, 53], Mathematical model (cancer) |
| $\theta_M$ | $M_1$ diff. upregulation by $K_A$ | $5.0 \times 10^{-8}$ day$^{-1}$ | [53], Mathematical model (cancer) |
| $\theta_K$ | $K_0$ activation by $M_1$ | 0.2 day$^{-1}$ | See Supplementary Section S2.3 |
| $\gamma$ | Proportion of $K_A$ relative to $M_1$ $E$ cell lysis | 0.8 (unitless) | Guess |
| $\omega$ | Lysis rate of E by immune cells/mL | $10^{-5}$ cells$^{-1}$ $\cdot$ mL $\cdot$ day$^{-1}$ | [53], Mathematical model (cancer) |
| $\rho_0$ | Proportion of $E_0$ cleared retrograde | 0.1 (unitless) | Guess |
| $\rho_F$ | Attachment rate of $E_F$ to $E_A$ | 0.1 day$^{-1}$ | Guess |
| $a_E$ | Endometrial shedding coefficient | $2 \times 10^4$ cells $\cdot$ mL$^{-1}$ $\cdot$ day$^{-1}$ | See Supplementary Supplementary S2.1 |
| $b_E$ | Endometrial shedding shape control | 160 | See Supplementary Supplementary S2.1 |
| $d_E$ | Endometrial shedding phase alignment | 12 days | See Supplementary Supplementary S2.1 |
| $\bar{\mu}_E$ | Surrogate model $E$ influx | 1260 cells $\cdot$ mL$^{-1}$ $\cdot$ day$^{-1}$ | Avg. cyclic (Eq. (9)) |

**Table S1:** Parameter values used in the model. Due to a lack of available data on the immune system response to endometriosis, many parameters are taken from literature on immune response to cancer or infection. This assumes that the timescales of the immune cell responses to endometrial cells are similar to those in cancer and infections. Further details on parameterisation are given in Supplementary S2.

| State | Initial Count (cells/mL) |
|---|---|
| $M_0(0)$ | $9.0 \times 10^5$ |
| $M_1(0)$ | 100 |
| $M_2(0)$ | $4.5 \times 10^4$ |
| $K_0(0)$ | $5.0 \times 10^5$ |
| $K_A(0)$ | $2.0 \times 10^4$ |
| $E_0(0),\ E_F(0),\ E_A(0)$ | 0 |

**Table S2:** Initial conditions. The total macrophage concentration $(M_0 + M_1 + M_2)$ was estimated from [5, 56] and the values for each state were then calculated using steady state equations. The resting natural killer cell concentration $(K_0)$ was taken from the control data in [25], and the activated natural killer was determined from the resulting steady state.

### S2 Parameterisation

#### S2.1 Eutopic endometrial cells

We fit a function for the eutopic endometrial cells using data on menstrual blood volume [52] and concentration of live stromal and epithelial cells found in uterine menstrual blood [13]. We assume the following function shape for $E_0$, as given in Eq. (9):

$$\mu_E(t) = \alpha \left[ \sin\left( \frac{t+d}{28}\pi \right) \right]^b . \tag{S1}$$

We determine the shape parameters by normalising the average volumes of menstrual blood across the four groups considered in [52]. We choose the shape parameters to be: $b = 160$ and $d = 12$. A plot of the $E_0$ timeseries using these shape parameters in comparison to the normalised data from [52] is shown in Fig. S1.

In order to choose a value for the coefficient $\alpha$, we assume the peak value of the $E_0$ is approximately $10^4$ cells/mL. A study on stem and progenitor cells in uterine menstrual fluid, collected through aspiration of the uterine cavity via the cervix, measured live cell concentrations of $3.8 \times 10^4$ and $0.8 \times 10^4$ cells/mL for mesenchymal stem cells and epithelial progenitor cells respectively [13]. It is important to note that concentration here is per uterine fluid, which would be diluted when mixed with the peritoneal fluid. We choose a coefficient of $\alpha = 2 \times 10^4$ cells/mL which results in a peak concentration of $\hat{E}_0 = 1.4 \times 10^4$ cells/mL.

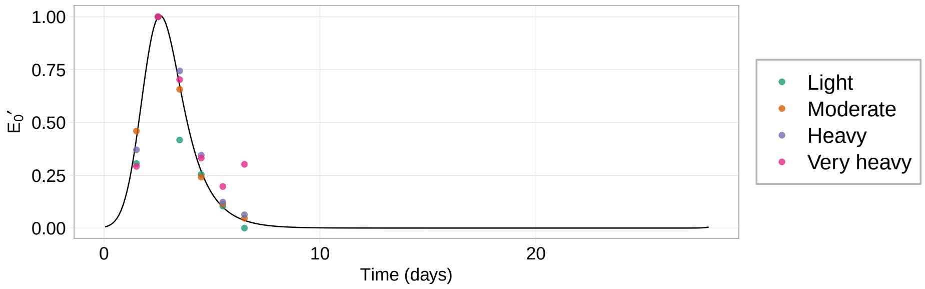


**Figure S1:** The solution for normalised $E_0$, $E_0'$, with the chosen shape parameters compared to menstrual blood volume data from [52], which we have normalised to peak volume in each group. Here $\alpha' = 1.45$, $b = 160$, $d = 12$, and $\delta_{E_0} = 1$ day$^{-1}$.

#### S2.2 Macrophage

Macrophage influx, capacity, and clearance are chosen to produce a total macrophage profile similar to those observed clinically, assuming the total concentration of macrophages does not change. Studies have shown that macrophage counts may increase in endometriosis [15, 58], however the peritoneal fluid volume may also increase [57]. Consequently, we can assume that that the average concentration is approximately constant, which is supported by many studies [5, 7, 56].

The steady state equation for the total macrophage in the system is:

$$\frac{dM_T}{dt} = \mu_M + \eta_M \left(1 - \frac{M_T}{M_C}\right) M_T - \delta_M M_T = 0 \tag{S2}$$

Assuming $\eta_M = 0.2$ day$^{-1}$, $M_C = 10^6$ cells/mL, and $\delta_M = 0.02$ day$^{-1}$ [19, 20], and $M_T \approx 10^6$ cells/mL [5, 56], we estimate $\mu_M = 10^4$ cells $\cdot$ mL$^{-1}$ $\cdot$ day$^{-1}$.

We can determine the steady state levels of macrophage states, with negligible endometrial cells and $K_A$ concentration from Table S3, to be:

$$M_0 \approx 9 \times 10^5; M_1 \approx 10^2; M_2 \approx 4 \times 10^4 \text{ cells/mL}. \tag{S3}$$

#### S2.3 Natural killer cells

We estimate the NK cell equation parameters using data from the literature on levels of natural killer and macropahge cells in peritoneal fluid (PF). To do this, we use control participant data from studies on PF in endometriosis. We make the assumption that in these control systems the amount of endometrial cells present in

| Cell type/state | Literature Data | Estimated concentration when $E_F + E_A \approx 0$ |
|---|---|---|
| Total natural killer | Ratio NK:M=10:57 [6] | $K_0 + K_A = 2 \times 10^5$ cells/mL |
| Activated natural killer | $K_A/(K_0 + K_A) = 0.15$ [17] | $K_A = 0.3 \times 10^5$ cells/mL |

**Table S3:** Estimated concentrations of natural killer cells states based on data from the literature. Note, this assumes a total macrophage concentration of $M_0 + M_1 + M_2 \approx 9.5 \times 10^5$ cells/mL, as estimated from the steady state equations for the total macrophage ($M_0 + M_1 + M_2$) system.

the system is negligible. Table S3 below summarises the data from the literature on NK concentrations in peritoneal fluid.

Under this condition, we get the following steady state equations for our natural killer cells:

$$\begin{aligned} \frac{dK_A}{dt} &= \theta_K K_0 \left( \frac{M_1}{C_{M_1} + M_1} \right) + \eta_K \left( 1 - \frac{K_A + K_0}{K_C} \right) K_A - \delta_K K_A = 0, \\ \frac{d(K_0 + K_A)}{dt} &= \mu_K + \eta_K \left( 1 - \frac{K_0 + K_A}{K_C} \right) K_A - \delta_K (K_0 + K_A) = 0. \end{aligned} \tag{S4}$$

We assume $\delta_K = 0.02$ day$^{-1}$ [55] (estimated from macaque data); and NK carrying capacity similar to M, $K_C = M_C = 10^6$ cells/mL. In [55], NK proliferation rate in macaques, assuming a linear form of the proliferation term, was approximately 0.01 day$^{-1}$ (influx not stated). We choose a value of $\eta_K = 0.02$ day$^{-1}$, which gives $\theta_K \approx 0.7$ and $\mu_K \approx 3.5 \times 10^3$ cells $\cdot$ mL$^{-1}$ $\cdot$ day$^{-1}$ using the macrophage steady state values from Supplementary S2.2.

### S3 Steady state analysis

To understand how the system is sensitive to system parameters, we suppose that there exists steady states of the following descriptions:

1. Endometrial free and immune free (the trivial steady state),
2. Endometrial free, natural killer cell free and macrophage present,
3. Endometrial free, macrophage free and natural killer cell present,
4. Endometrial free and immune present.

Here, we will discuss each of the above steady states. We first define the Jacobian of the system, $\frac{dX_i}{dt} = f_i(\mathbf{X}, \mathbf{\Theta})$, for $i = 1, ..., 8$, as:

$$J(\mathbf{X}, \mathbf{\Theta}) := \left[\frac{\partial \mathbf{f}}{\partial X_1}, \ldots, \frac{\partial \mathbf{f}}{\partial X_n}\right]^T,$$

where $\mathbf{X} = (M_0, M_1, M_2, K_0, K_A, E_0, E_F, E_A)$ is the system state variable, and $\mathbf{\Theta}$ contains the parameters of the system. For a given steady state, $\mathbf{X}^*$, we can then determine its stability via considering the eigenvalues of the system:

$$|J - \lambda I|_{\mathbf{X}^*} = 0.$$

A disease free state is one where $E_A^* = 0$. Furthermore, we know that in the disease free state that immune cells are present. Therefore, we are interested in knowing how we can modify the system parameters to ensure that the disease free state is always stable.

#### Case 1: Endometrial free and immune free (the trivial steady state)

We consider the trivial endometrial free and immune free steady state, i.e. $E_0 > 0$, $E_F = 0$, $E_A = 0$, $M_0 = 0$, $M_1 = 0$, $M_2 = 0$, $K_0 = 0$, and $K_A = 0$. The linear stability analysis gives:

$$\begin{aligned}
0 =& |J - \lambda I|_{\mathbf{X}^*}, \\
0 =& (\lambda + \delta_M - \mu_M)(\lambda + \beta_{12} + \beta_{21} + \delta_M)(\lambda + \beta_2 + \delta_M) \\
&\times (\lambda + \delta_K)(\lambda + \delta_K - \eta_K) \\
&\times (\lambda + \delta_{E_0})(\lambda + \delta_E + \rho_F)(\lambda + \delta_E),
\end{aligned}$$

which is always stable, for $\delta_K > \eta_K$ (which are the rates of activated natural killer cell removal and proliferation) and $\delta_M > \mu_M$ (which are the rates of macrophage removal and proliferation) since $\mathbf{\Theta} > \mathbf{0}$.

#### Case 2: Endometrial free, natural killer cell free and macrophage present

We consider the endometrial free, natural killer cell free and macrophage present steady state. Here $E_0 > 0$, $E_F = 0$, $E_A = 0$, $K_0 = 0$, $K_A = 0$, $M_0 > 0$, $M_1 > 0$, and $M_2 > 0$. The linear stability analysis gives:

$$\begin{aligned}
0 =& |J - \lambda I|_{\mathbf{X}^*}, \\
0 =& \frac{-1}{k_{M_2} + M_2}(\lambda + \delta_M - \mu_M)(\lambda + \beta_{12} + \beta_{21} + \delta_M)(\lambda + \beta_2 + \delta_M) \\
&\times \left(\lambda + \frac{\theta_K M_1}{C_{M_1} + M_1} + \delta_K\right)(\lambda + \delta_K - \eta_K) \\
&\times (\lambda + \delta_{E_0})(\lambda + (1-\gamma)M_1\omega + \delta_E + \rho_F)\left(\lambda + (1-\gamma)M_1\omega + \delta_E - \frac{\eta_A M_2}{C_{M_2} + M_2}\right),
\end{aligned}$$

which is stable for:

$$(1-\gamma)M_1\omega + \delta_E > \frac{\eta_A M_2}{C_{M_2} + M_2},$$

in addition to the cases stated above. We interpret this inequality as a requirement for the rate of attached endometrial cell removal to be greater than the rate of attached endometrial cell proliferation.

**Case 3: Endometrial free, macrophage free and natural killer cell present**

We consider the endometrial free, macrophage free and natural killer cell present steady state. Here $E_0 > 0$, $E_F = 0$, $E_A = 0$, $M_0 = 0$, $M_1 = 0$, $M_2 = 0$, $K_0 > 0$, and $K_A > 0$. The linear stability analysis gives:

$$\begin{aligned}
0 =& |J - \lambda I|_{\mathbf{X}^*}, \\
0 =& (\lambda + \delta_M - \mu_M)(\lambda + \beta_{12} + \beta_{21} + \delta_M)\left(\lambda + \beta_2 + \frac{\theta_M K_A}{C_{K_A} + K_A} + \delta_M\right) \\
&\times (\lambda + \delta_K)\left(\lambda + \delta_K - \eta_K\left(1 - \frac{K_0 + 2K_A}{K_C}\right)\right) \\
&\times (\lambda + \delta_{E_0})(\lambda + \gamma K_A \omega + \delta_E + \rho_F)(\lambda + \gamma K_A \omega + \delta_E),
\end{aligned}$$

which is stable for:

$$\delta_K > \eta_K\left(1 - \frac{K_0 + 2K_A}{K_C}\right),$$

in addition to the balance of macrophage removal and supply.

**Case 4: Endometrial free and immune present**

We lastly consider the endometrial free and immune present steady state, which represents the healthy system. Here $E_0 > 0$, $E_F = 0$, $E_A = 0$, $M_0 > 0$, $M_1 > 0$, $M_2 > 0$, $K_0 > 0$, and $K_A > 0$, which can be considered as a combination of the two previously mention states. Therefore, since both of those states are stable, we also expect this state to also exhibit similar stability. The linear stability analysis gives:

$$\begin{aligned}
0 =& |J - \lambda I|_{\mathbf{X}^*}, \\
0 =& \frac{1}{k_{M_2} + M_2}\left(\lambda + \delta_M - \mu_M \frac{2(M_0 + M_1 + M_2) - M_C}{M_C}\right) \\
&\times \left\{\left(\frac{\theta_M C_{K_A} M_0}{C_{K_A} + K_A}\right)\left(\frac{\theta_K K_0 C_{M_1}}{C_{M_1} + M_1}\right)(\lambda + \beta_2 + \beta_{21} + \delta_M)\right. \\
&\left. - (\lambda + \beta_{12} + \beta_{21} + \delta_M)\left(\lambda + \beta_2 + \frac{\theta_M K_A}{C_{K_A} + K_A} + \delta_M\right)\left(\lambda + \delta_K - \eta_K\left(1 - \frac{K_0 + 2K_A}{K_C}\right)\right)\right\} \\
&\times (\lambda + \delta_{E_0})(\lambda + [(1-\gamma)M_1 + \gamma K_A]\omega + \delta_E + \rho_F)\left(\lambda + [(1-\gamma)M_1 + \gamma K_A]\omega + \delta_E - \frac{\eta_A M_2}{C_{M_2} + M_2}\right).
\end{aligned}$$

The first eigenvalue is associated with the resting macrophage removal and proliferation, and we therefore expect this state to exhibit stability when

$$\delta_M > \mu_M \frac{2(M_0 + M_1 + M_2) - M_C}{M_C}.$$

The second, third, and forth eigenvalues (the terms in the curly brackets) are associated with the activation, removal, and polarisation between anti- and pro-inflammatory type macrophage, and also the activation and removal of activated natural killer cells. However, these eigenvalues are present as a cubic polynomial, which cannot be solved simply. We therefore hypothesise on it's stability from the stablility of the natural killer free and the macrophage free states respectively, and assume that these eigenvalues are also stable.

The fifth eigenvalue is associated with the removal of eutopic endometrial cells, and is stable since $\delta_{E_0} > 0$. The sixth eigenvalue is associated with the removal and attachment of in fluid endometrial cells, which is stable, since

$$[(1-\gamma)M_1 + \gamma K_A]\omega + \delta_E + \rho_F > 0.$$

The seventh and last eigenvalue is associated with the removal and proliferation of attached endometrial cells, which is stable when

$$[(1-\gamma)M_1 + \gamma K_A]\omega + \delta_E > \frac{\eta_A M_2}{C_{M_2} + M_2}, \tag{S5}$$

i.e. when the rate of attached endometrial cell removal is greater than the rate of attached endometrial cell proliferation. However, since this depends on the state of the anti- and pro-inflammatory macrophages, as well as the activated natural killer cells, this inequality will not always hold true. We therefore find that this is a necessary condition for the disease-free, endometrial free steady state to exist, otherwise, we hypothesise, the system will exhibit a diseased, endometrial present state.

### S4 System response with increased endometrial attachment rate

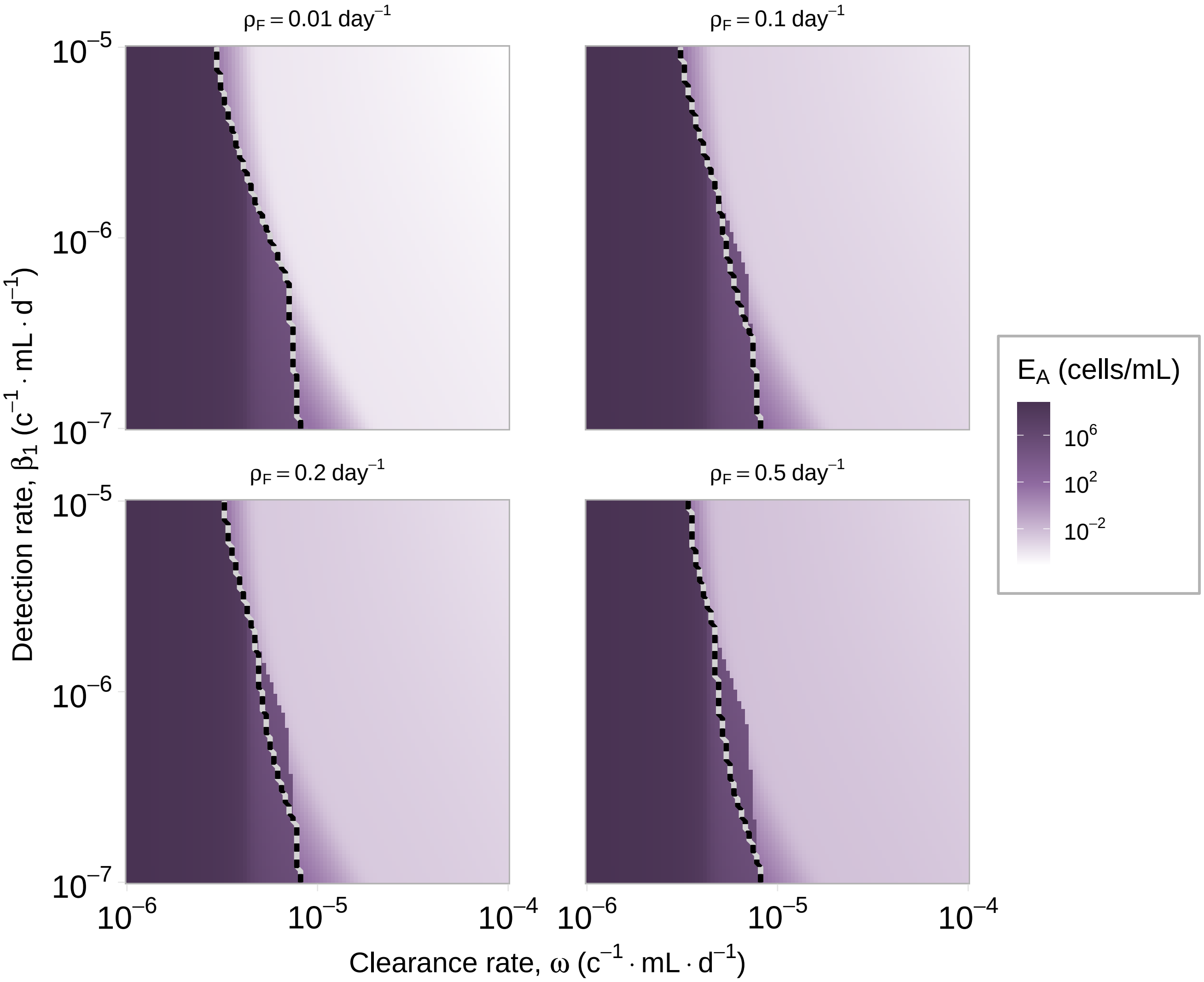


**Figure S2:** Heatmaps showing the level of attached endometrial cells at the end of a cycle with varying immune detection and clearance efficacy for different endometrial cell attachment rates ($\rho_F$). The attachment rate for the results in the main manuscript is $\rho_F = 0.1 \text{ day}^{-1}$. The dashed line indicates the partition between high and low attachment regions according to the inequality in Eq. (11). Units (c$^{-1}$ · mL · d$^{-1}$) are short for (cells$^{-1}$ · mL · day$^{-1}$). These results indicate that increasing the attachment rate of the endometrial cells ($\rho_F$) results in small increases to $E_A$ in the low disease state, but does change the key results of the paper.

### S5 Bifurcation diagrams

#### S5.1 Detection rate

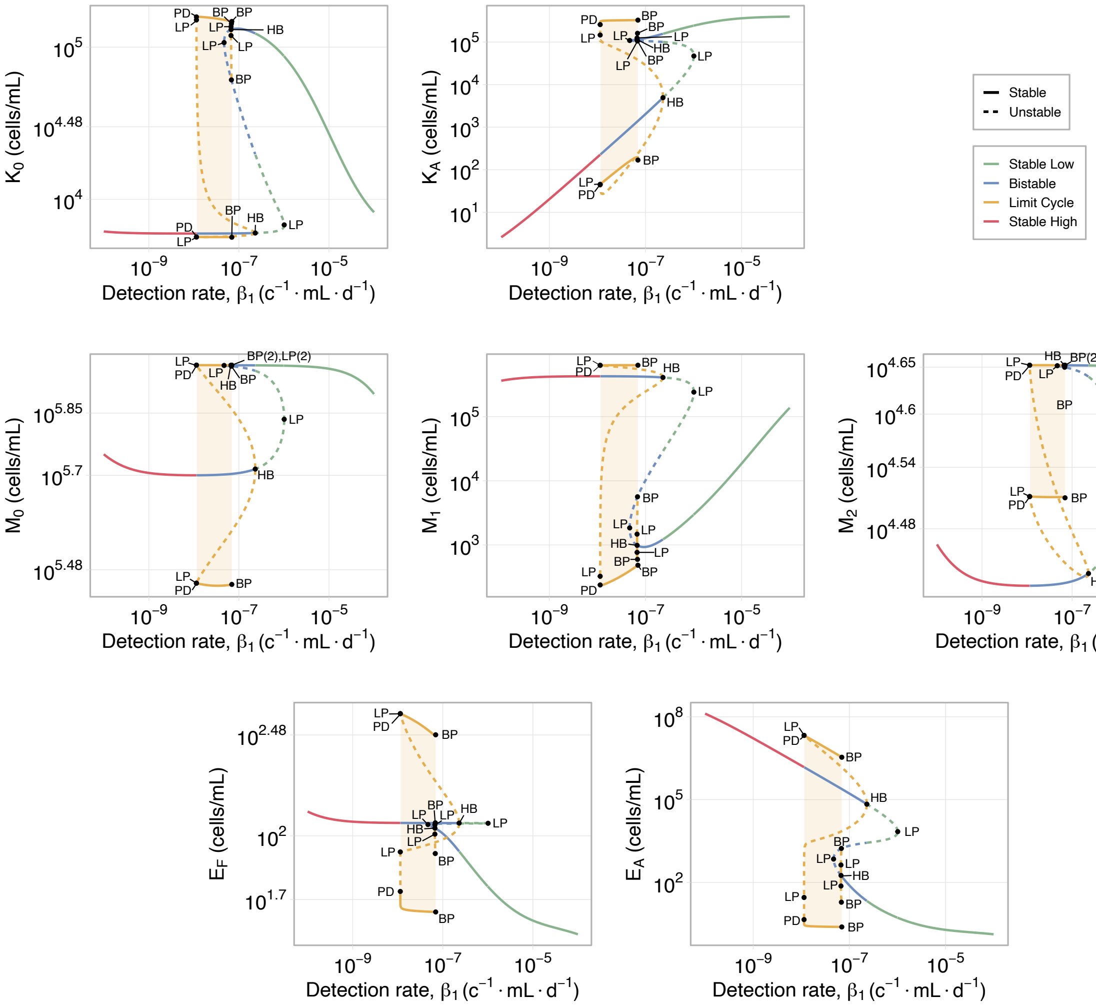


**Figure S3:** Bifurcation diagrams for the rate at which the endometrial cells are detected by the immune system (via macrophages), $\beta_1$. Clearance rate is fixed at $\omega = 10^{-5}$ cells$^{-1}\cdot$mL$\cdot$day$^{-1}$, and influx at $\rho_0 = 0.1$. Units (c$^{-1}\cdot$mL$\cdot$d$^{-1}$) are short for (cells$^{-1}\cdot$mL$\cdot$day$^{-1}$). Point labels are BP: bifurcation point, LP: limit point, PD: period doubling, HB: Hopf bifurcation. High (pink) is hypothesised to be the high disease state, low (green) is hypothesised to be the low or no disease state, and bistable (blue) is the transition region in Fig. 6(a). The bifurcation analysis was performed using AUTO-07p [60].

### S5.2 Clearance rate

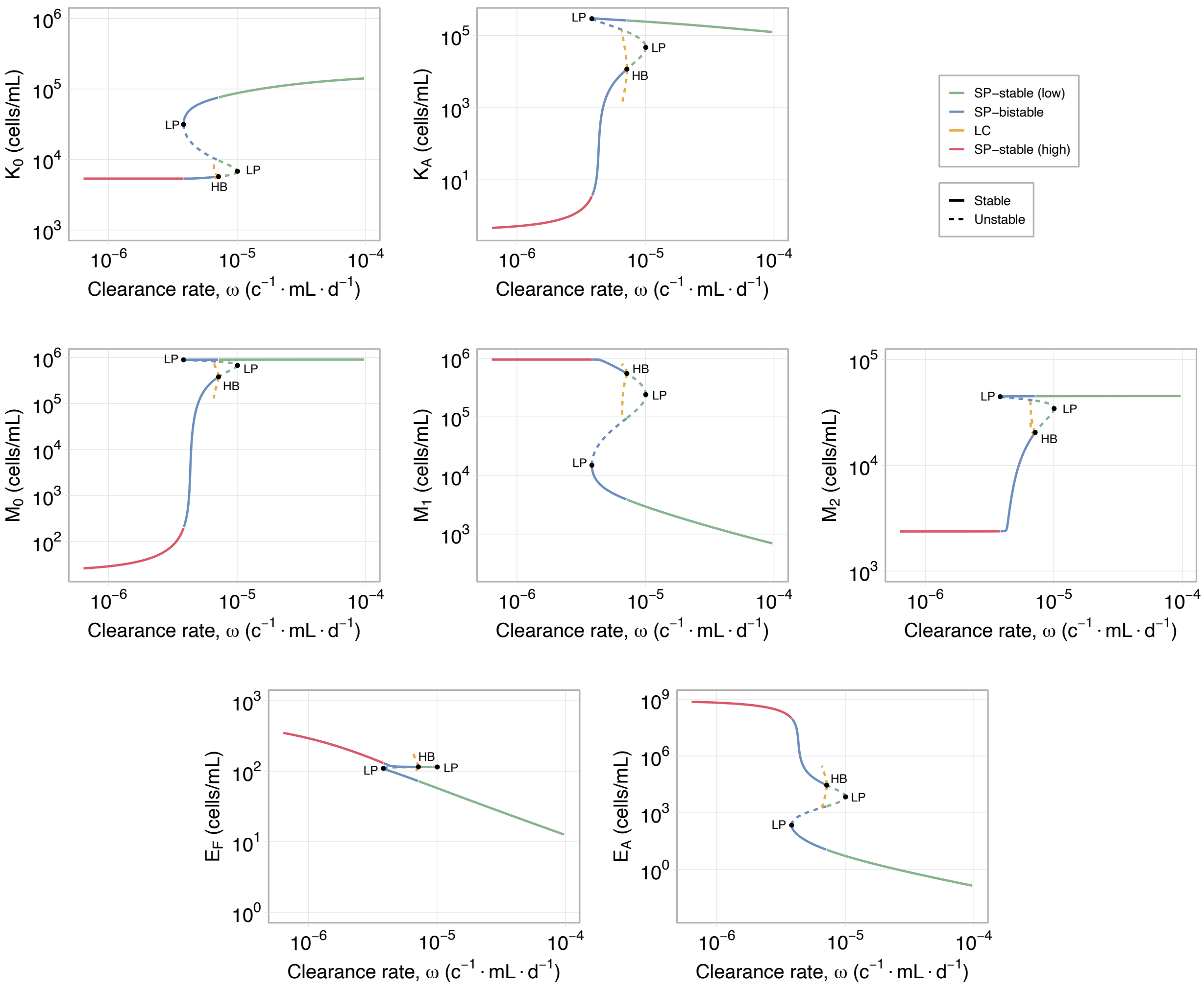


**Figure S4:** Bifurcation diagrams for the rate at which the endometrial cells are cleared by the immune system, $\omega$. Detection rate is fixed at $\beta_1 = 10^{-6}$ cells$^{-1}$ · mL · day$^{-1}$, and influx at $\rho_0 = 0.1$. Units (c$^{-1}$ · mL · d$^{-1}$) are short for (cells$^{-1}$ · mL · day$^{-1}$). Point labels are LP: limit point, HB: Hopf bifurcation. High (pink) is hypothesised to be the high disease state, low (green) is hypothesised to be the low or no disease state, and bistable (blue) is the transition region in Fig. 6(a). The bifurcation analysis was performed using AUTO-07p [60].

#### S5.3 Co-dimensional bifurcation diagram and examples

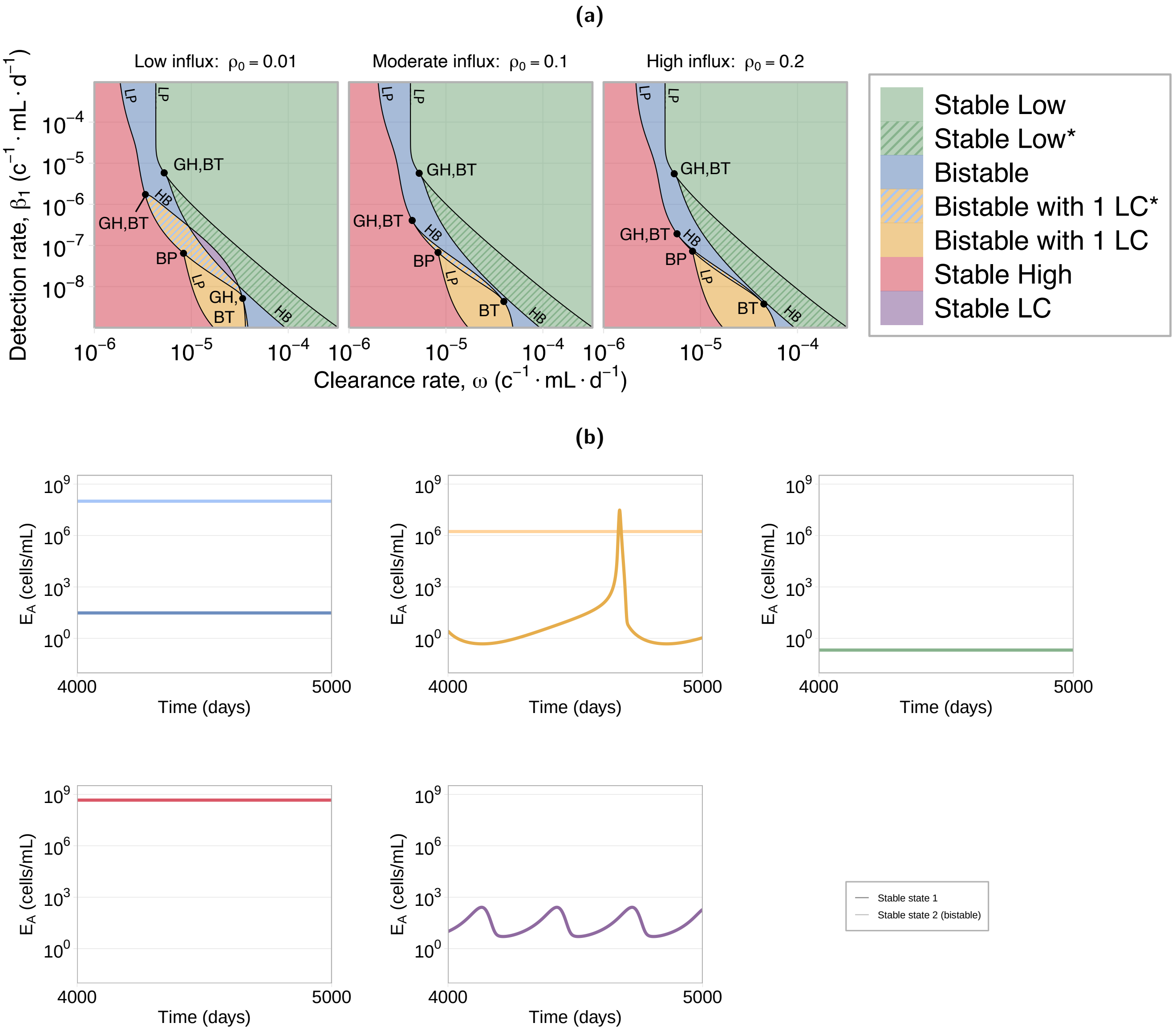


**Figure S5:** (a) Co-dimensional bifurcation, *Regions with additional unstable branches, LC: limit cycle. Units ($c^{-1} \cdot mL \cdot d^{-1}$) are short for ($cells^{-1} \cdot mL \cdot day^{-1}$). Branch labels are HB: Hopf bifurcation continuation, LP: Limit point continuation. Point labels are BP: bifurcation point, GH: generalised Hopf bifurcation, BT: Bogdanov-Takens bifurcation. Points labelled 'GH,BT' are GH and BT points that are located very close together. (b) Examples for each of the phases determined in the bifurcation analysis (using constant influx surrogate model). The stability regions found are: bistable (blue), bistable with a stable limit cycle (orange), stable low (green), stable high (red), and stable limit cycle (purple). The bifurcation analysis was performed using AUTO-07p [60].

### S6 Analysis of upregulation of M2-type macrophages by attached endometrial cells

#### S6.1 Modified system of equations

We modify the equations for $M_0$ and $M_2$ (Eqs. (1) and (3) respectively) to incorporate upregulation of M2-type macrophage activation in the presence of attached endometrial cells. However, we note that upregulation of $M_2$ by $E_A$ may be more associated with mid-late stage disease [30], where it is believed that the adaptive immune system will also play a crucial role. We modify the $M_2$ activation term as follows:

$$\begin{aligned}\frac{dM_0}{dt} &= \mu_M + \eta_M\left[1 - \left(\frac{M_0+M_1+M_2}{M_{\mathrm{C}}}\right)\right]\left[M_0+M_1+M_2\right] - \theta_M\left(\frac{K_A}{C_{K_A}+K_A}\right)M_0 \\ &\quad - \beta_1\left(E_F+E_A\right)M_0 - \underbrace{\beta_2\left(1+\alpha\frac{E_A}{C_{E_A}+E_A}\right)M_0}_{\text{Upregulated } M_2 \text{ activation due to } E_A} - \delta_{M_0}M_0,\end{aligned} \tag{S6}$$

$$\frac{dM_2}{dt} = \underbrace{\beta_2\left(1+\alpha\frac{E_A}{C_{E_A}+E_A}\right)M_0}_{\text{Upregulated } M_2 \text{ activation due to } E_A} + \beta_{12}M_1 - \beta_{21}M_2 - \delta_{M_2}M_2. \tag{S7}$$

where $\alpha$ is the effect of upregulation of the attached endometrial cells ($E_A$) and $C_{E_A}$ is the action limiting capacity of the $E_A$ cells. Here we use the value $C_{E_A} = 10^5$ cells/mL. When $\alpha = 0$ we recover our original equations for the macrophage dynamics, defined in Eqs. (1) to (3).

#### S6.2 Bifurcation analysis

We perform a one dimensional bifurcation analysis over the (unitless) parameter $\alpha$ to investigate the potential effect of upregulation on the system, shown in Figure S6. These results indicate that increasing the upregulation of $M_2$ by $E_A$ (i.e. $\alpha$), births a stable high disease state. The low disease state remains locally stable (solid green line), while the high disease state transitions from unstable (not shown), to stable (solid red line). This indicates that, when starting from the low disease state, simply increasing upregulation of $M_2$ by $E_A$ (i.e. $\alpha$) will not lead to disease onset (i.e. the high disease state), rather the system will be maintained in low disease. We therefore conclude that upregulation of $M_2$ by $E_A$ alone is not sufficient for disease onset.

Furthermore, these results display heterogeneity in the activation state of M2-type macrophages. Namely, we observe an increased level of $M_2$ that is associated with the high diseased state, noting that this high disease state requires an additional trigger for emergence to occur.

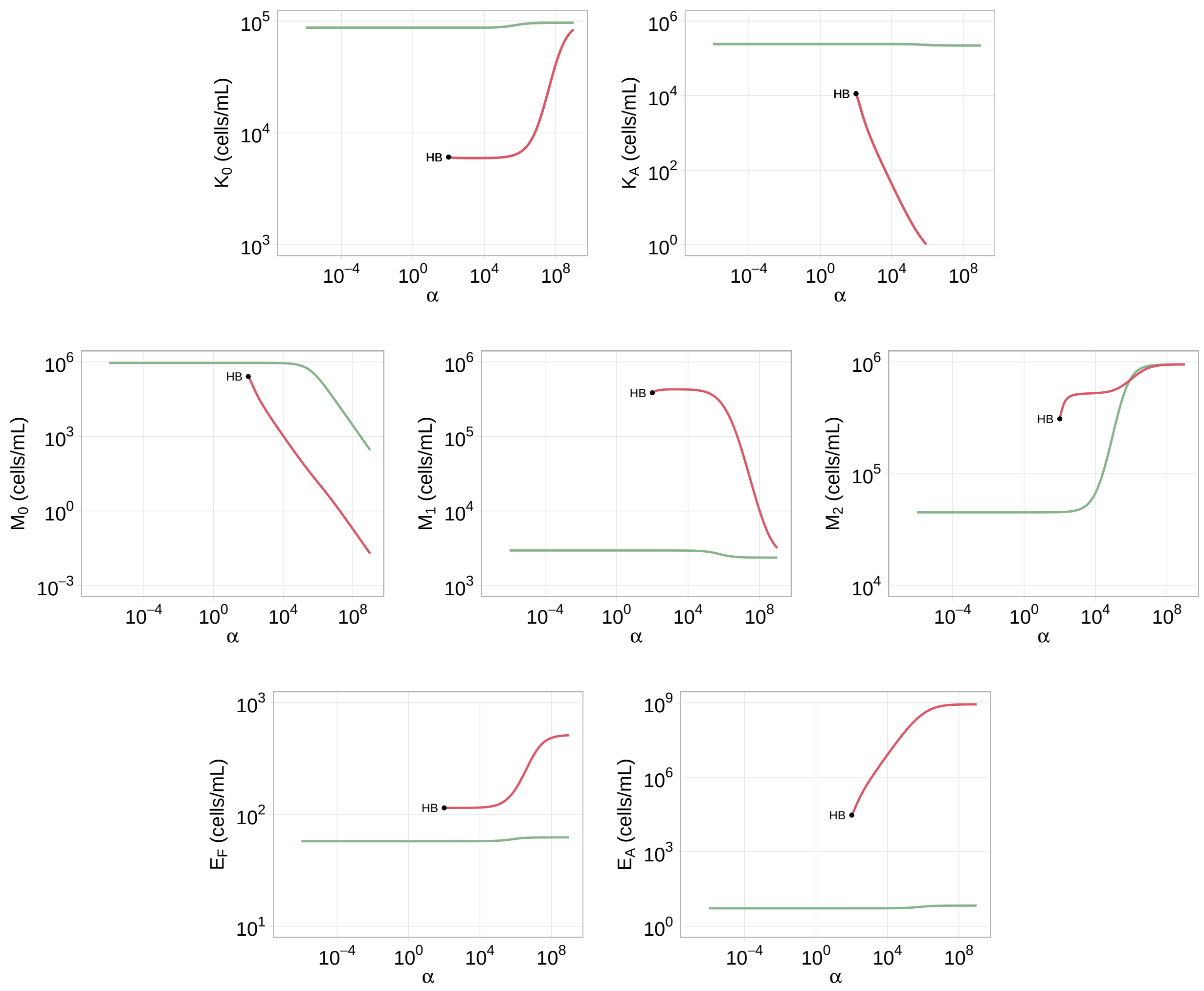


**Figure S6:** Bifurcation analysis over $\alpha$, the level of upregulation of $M_2$ activation by $E_A$. HB: Hopf Bifurcation point.
